# Molecular interactions underlying the phase separation of HP1α: Role of phosphorylation, ligand and nucleic acid binding

**DOI:** 10.1101/2022.06.20.496886

**Authors:** Cheenou Her, Tien M. Phan, Nina Jovic, Utkarsh Kapoor, Bryce E. Ackermann, Azamat Rizuan, Young Kim, Jeetain Mittal, Galia T. Debelouchina

## Abstract

Heterochromatin protein 1α (HP1α) is a crucial component for the proper maintenance of chromatin structure and function. It has been proposed that HP1α functions through liquid-liquid phase separation (LLPS), which allows it to sequester and compact chromatin into transcriptionally repressed heterochromatin regions. In vitro, HP1α can form phase separated liquid droplets upon phosphorylation of its N-terminus extension (NTE) and/or through interactions with DNA and chromatin. While it is known that LLPS requires homodimerization of HP1α and that it involves interactions between the positively charged hinge region of HP1α and the negatively charged phosphorylated NTE or nucleic acid, the precise molecular details of this process and its regulation are still unclear. Here, we combine computational modeling and experimental approaches to elucidate the phase separation properties of HP1α under phosphorylation-driven and DNA-driven LLPS conditions. We also tune these properties using peptides from four HP1α binding partners (Sgo1, CAF-1, LBR, and H3). In phosphorylation-driven LLPS, HP1α can exchange intradimer hinge-NTE interactions with interdimer contacts, which also leads to a structural change from a compacted to an extended HP1α dimer conformation. This process can be enhanced by the presence of positively charged peptide ligands such as Sgo1 and H3 and disrupted by the addition of negatively charged or neutral peptides such as LBR and CAF-1. In DNA-driven LLPS, both positively and negatively charged peptide ligands can perturb phase separation. Our findings demonstrate the importance of electrostatic interactions in the LLPS of HP1α where binding partners can modulate the overall charge of the droplets and screen or enhance hinge region interactions through specific and non-specific effects. Our study illuminates the complex molecular framework that can fine tune the properties of HP1α and that can contribute to heterochromatin regulation and function.

## Introduction

Heterochromatin is a fundamental architectural feature of eukaryotic chromosomes that has essential roles in processes such as gene repression and silencing, chromosome segregation, and DNA repair.^1–6^ The formation and dissolution of heterochromatin allow for transcriptional control where genes can be silenced or upregulated, resulting in distinct cellular phenotypes encoded by the same genome.^3,7,8^ Heterochromatin function and organization is partially attributed to the ability of heterochromatin protein 1 (HP1) to recruit ligands and spread across the genome.^9–12^ In humans, HP1 exists as three paralogs, HP1α, HP1β, and HP1γ, all of which have gene silencing roles, and two (HP1β and HP1γ) have been implicated in gene activation.^11^ Studies have found that changes in the expression levels of HP1 are linked to the progression of many forms of cancer. For example, reduced levels of HP1α have been associated with breast,^13,14^ brain,^15^ and colon cancer,^16,17^ while lowering the levels of HP1γ has been linked to ovarian cancer.^18^

The multi-functionality of HP1 proteins can be attributed to their structural complexity that enables a vast interaction network with DNA, RNA, and a wide array of nuclear proteins.^19–21^ They are multi-domain proteins that consist of three disordered regions, the N-terminal extension (NTE), the hinge region, and the C-terminal extension (CTE), along with two highly conserved folded domains, the chromodomain (CD) and the chromoshadow domain (CSD), which are topologically connected as shown in **Fig. 1a,b**. The presence of multiple domains allows HP1 paralogs to establish a complex interaction network with themselves and with other nuclear components. Interactions between the CSD domains are responsible for homodimerization which provides a hydrophobic binding surface for ligands that contain a PXVXL motif (where X denotes any amino acid).^11,22–24^ These interactions are responsible for the recruitment of additional proteins to heterochromatin. On the other hand, the CD recognizes and directly interacts with methylated lysine 9 on the histone H3 tail (H3K9me), an epigenetic mark associated with transcriptional repression.^25,26^ Compared with the CD and CSD, the flexible disordered regions are less conserved and may be responsible for the unique functional properties of different HP1 paralogs. The hinge region, which has patches of positively charged residues, can bind non-specifically to DNA and RNA,^27,28^ and these interactions have been implicated in heterochromatin maintenance.^29^ In addition, the interactions between the hinge region and the CTE are proposed to mediate an auto-inhibited dimer conformation.^30,31^

**Figure 1.**
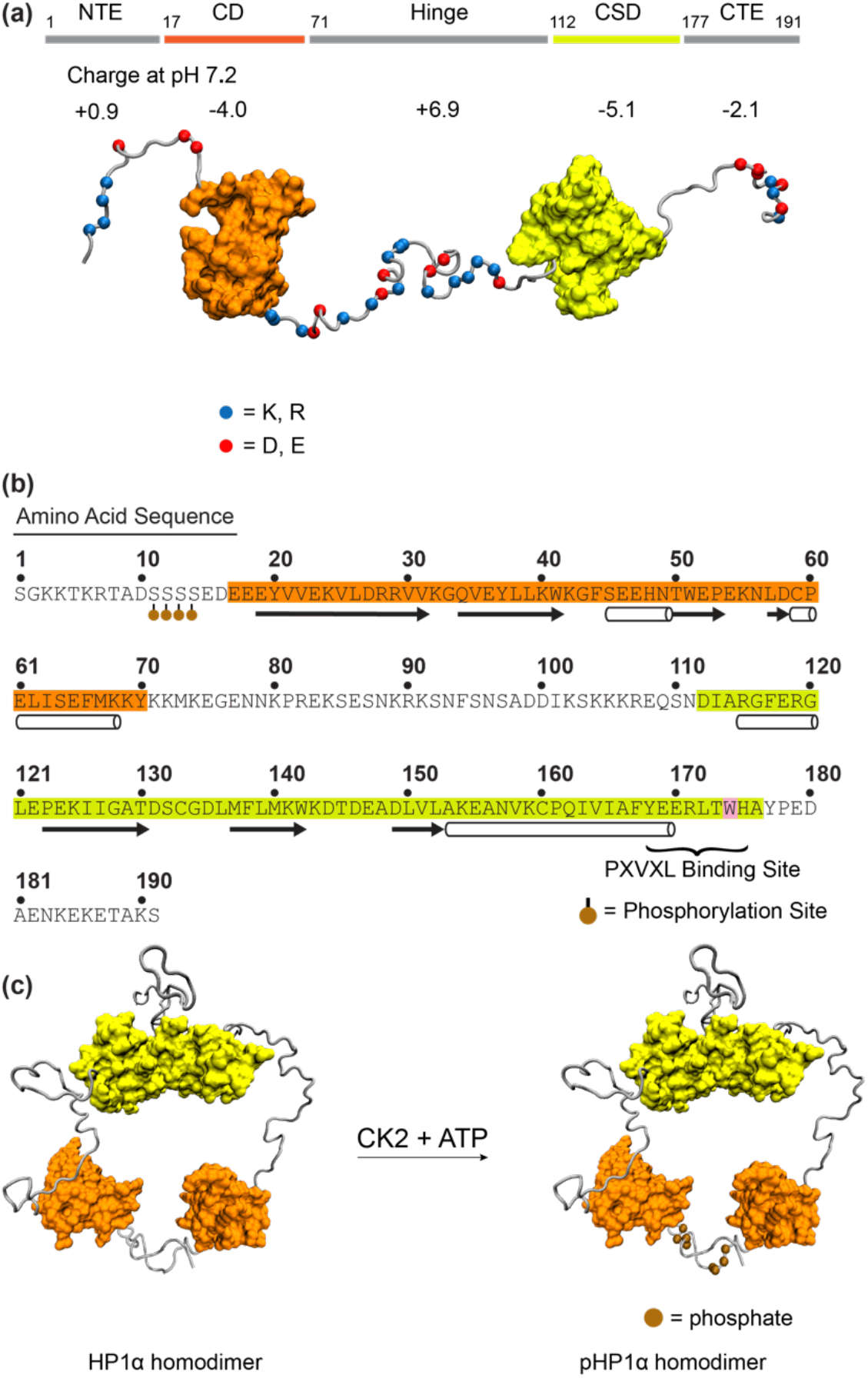
The properties of human heterochromatin protein 1α (HP1α). **(a)** A model depicting the structure of HP1α, comprising the N-terminus extension (NTE), the chromodomain (CD, orange), the hinge region, the chromoshadow domain (CSD, yellow), and the C-terminus extension (CTE). The charge at pH 7.2 is indicated for each region. **(b)** The amino acid sequence of HP1α with the CD highlighted in orange and the CSD highlighted in yellow. Trp 174 is highlighted in pink. The phosphorylation sites are shown at Ser 11, 12, 13, and 14. HP1α residues that bind the PXVXL/PXVXL-like motifs are indicated on the amino acid sequence. The secondary structures are based on the crystal structures with PBD code 3fdt for the CD and 3i3c for the CSD, respectively. **(c)** Schematic of the phosphorylation of HP1α using Casein Kinase II (CK2).

Recently, liquid-liquid phase separation (LLPS) was proposed as a mechanism for the arrangement of heterochromatin. In particular, HP1α, but not HP1β or HP1γ, has been shown to form liquid-like droplets at high protein concentration and low salt conditions.^27,32^ In addition, phosphorylation of the NTE or mixing with DNA promotes HP1α condensation. Interestingly, specific ligands, which directly interact with the CSD-CSD interface, can also enhance or attenuate HP1α phase separation.^31^ It has been suggested that phosphorylation or DNA-binding relieves autoinhibition and opens up the HP1α dimer, which provides an opportunity for multivalent interactions with other dimers and the formation of higher order oligomers.^31^ While this model suggests a central role for the open state in promoting multivalent interactions that drive LLPS, the precise molecular details are not well understood. For example, how do intra- and inter-dimer interactions change upon phosphorylation or in the presence of DNA binding, and how do these changes drive LLPS? And how do ligands tune these interactions to promote or inhibit phase separation? Uncovering the underlying forces that drive HP1α LLPS is vital to understanding the regulatory function of heterochromatin.

Here, we use a combination of computational and experimental techniques to describe the interaction network of HP1α in the context of NTE phosphorylation and binding with DNA. We probe the phosphorylation-induced conformational changes of the HP1α homodimer and characterize the molecular interactions underlying phosphorylation- or DNA-driven LLPS in the presence or absence of ligands. Our results paint a complex picture where HP1α must balance favorable hinge-phosphorylated NTE or hinge-DNA interactions with competing electrostatic interactions from the ligands. The findings of this study present a molecular framework for conceptualizing the mechanism and regulation of HP1α LLPS to understand heterochromatin formation and its functional relevance.

## Results

### Conformational changes of HP1α upon phosphorylation

HP1α typically functions as a homodimer formed by a CSD-CSD interaction interface with an estimated dissociation constant (K_d_) in the low nanomolar range^20^ (**Fig. S1**). The hinge region is enriched in lysine and arginine residues and is positively charged, while the CD, CSD, and CTE domains have an overall negative charge. The NTE region contains a stretch of four consecutive serine residues that can be phosphorylated *in vivo* by Casein Kinase II (CK2)^33^ (**Fig. 1c**). CK2 can also specifically phosphorylate these serine residues *in vitro*,^31,33^ producing up to four NTE phosphorylation sites as shown by LC-MS and MS-MS analysis (**Fig. S2 & Table S1**). Previous studies have shown that phosphorylation can promote the ability of HP1α to undergo LLPS.^31^ In our samples, HP1α phosphorylated *in vitro* (pHP1α) readily forms droplets at concentrations above 50 μM while no phase separation is observed for the wild-type protein up to 250 μM (**Fig. 2a**). It should be noted that at higher salt concentrations (e.g. 300 mM KCl), no liquid droplet formation is observed for either protein, which highlights the role of electrostatic attractions in promoting phase separation of pHP1α (**Fig. S3a**). Samples prepared at high salt will therefore serve as a no LLPS control in the phosphorylation-driven LLPS experiments described below.

**Figure 2.**
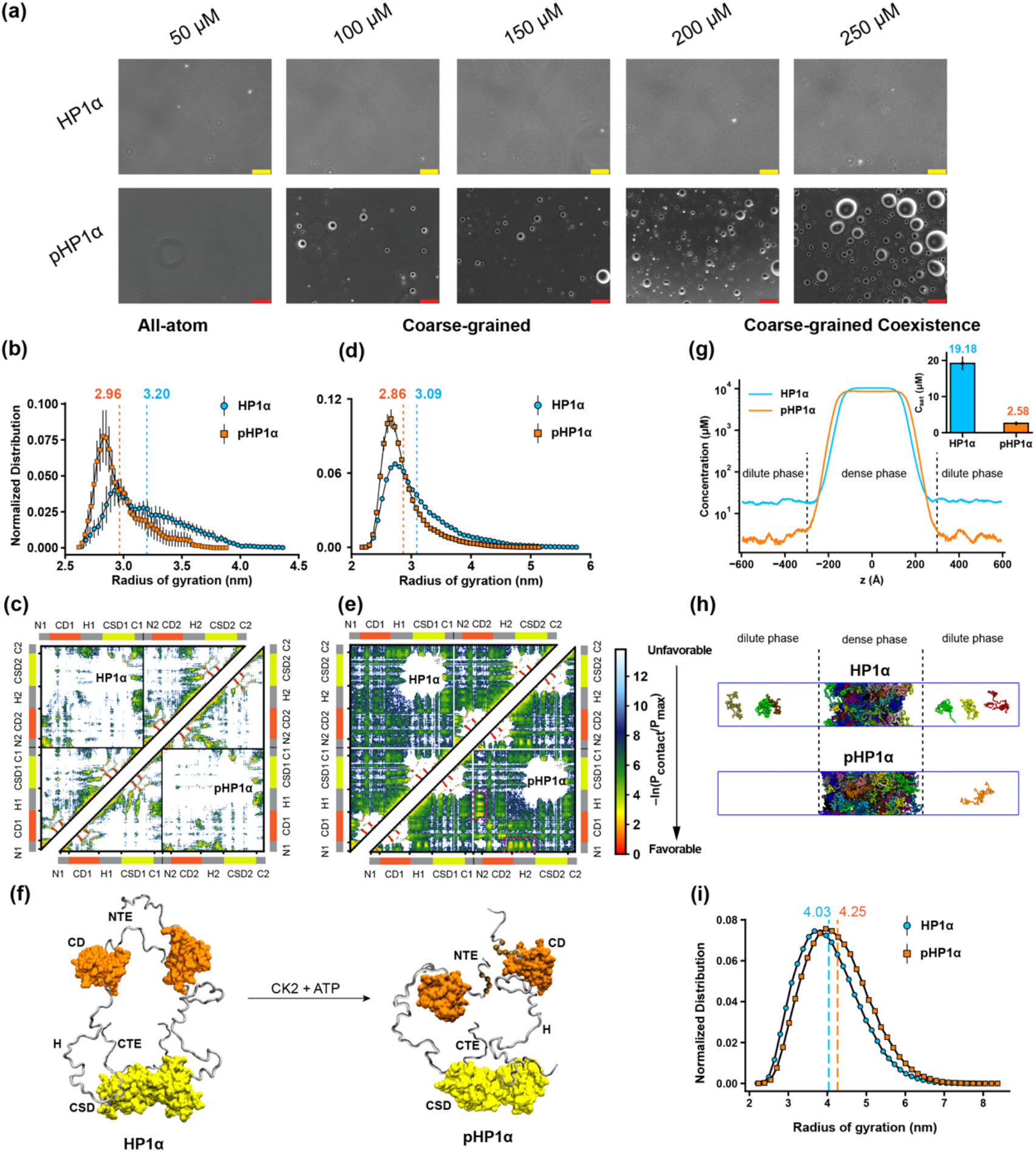
Effect of phosphorylation on the conformation and phase behavior of HP1α. **(a)** Brightfield microscopy images of HP1α and pHP1α with increasing protein concentration. The yellow and red scale bars represent 50 and 100 μm, respectively. **(b, d)** Rg distributions of HP1α and pHP1α homodimers in the AA and CG simulations. Dashed lines represent mean values of each distribution. **(c, e)** Contact maps of HP1α (top triangle) and pHP1α (bottom triangle), in AA and CG simulations. The intramolecular interactions within one chain and the intermolecular interactions between two chains are shown in the two small triangles and the off-diagonal quadrant, respectively. (**f**) Cartoons show the phosphorylation-induced conformation (randomly selected) of HP1α. **(g)** Density profiles and saturation concentrations (inset) of HP1α and pHP1α in coarse-grained coexistence simulations. (**h**) Snapshots of the condensates in the CG coexistence simulations. **(i)** The Rg distributions of HP1α and pHP1α in CG coexistence simulations. Errors of the Rg distributions are estimated using block averages with five blocks. The density profiles for HP1α and pHP1α in (g) are averaged over three replicas.

Previous literature has suggested that phosphorylation of the NTE can lead to an extended, more open conformation of the HP1α homodimer.^31^ This conformation can then promote attractive electrostatic interactions between the negatively charged NTE of one homodimer and the positively charged hinge region of another dimer, thus facilitating the LLPS process. The precise molecular details of how phosphorylation may change intramolecular (within the dimer) and intermolecular interactions between HP1α dimers are not entirely clear. For example, what are the dominant interactions before phosphorylation and how does modification alter the interaction landscape? And are there additional interactions beyond the hinge and the NTE that help drive oligomer formation? To address this, we performed molecular dynamics (MD) simulations using a current state-of-the-art all-atom (AA) protein model (Amber99SBws-STQ)^34^ with explicit solvent (TIP4P/2005).^35^ Full-length HP1α was constructed using PDB structural models 3fdt^36^ and 3i3c^37^ for the CD and CSD domains, respectively. The disordered regions (NTE, hinge, and CTE) were connected to the folded domains using MODELLER.^38^ To form the expected HP1α dimer configuration, the CSD domains were arranged to interact through the α-helix binding interface (3i3c structural model). The HP1α and pHP1α dimers were simulated for 5 μs using OpenMM (see Methods for details). Within the simulation time, the proteins sample a wide variety of configurations as shown in **Movies S1** and **S2**.

To gain insight into the phosphorylation-induced changes in the protein structural properties, we calculated the radius of gyration (R_g_) distributions of HP1α and pHP1α homodimers. Contrary to expectations based on previous work,^31^ we find that the pHP1α conformations (R_g_ = 2.96 ± 0.21 nm) are more compact than the HP1α structures (R_g_ = 3.20 ± 0.31 nm) (**Fig. 2b**). The HP1α dimer has a net negative charge (q = -5.8) and phosphorylation makes it even more negative (q = -21.8), which should lead to chain expansion unless segregation of like-charges in specific regions can promote interactions between oppositely charged residues.^39,40^ Indeed, phosphorylation concentrates negative charge on the NTE, which can now interact with the positively charged hinge region. To characterize molecular interactions within the HP1α dimer and changes upon phosphorylation causing the observed conformational differences, we computed the number of intra- (within an HP1α monomer) and intermolecular (between the two HP1α monomers) van der Waals (vdW) contacts formed by each residue from the all-atom simulation data (**Fig. 2c**). The contact map is relatively sparse given the limited sampling over the 5 μs simulation time, but a few critical observations can be made. In addition to contacts formed within the folded CD and CSD domains, NTEs (N1:N2) interact strongly with each other, presumably due to charge attraction between K/R and D/E residues, in both HP1α and pHP1α. Local interactions between folded CDs and CSDs with disordered segments (NTE and CTE) are also present in HP1α. We also observe inter- and intramolecular interactions between the CTE and the hinge in both HP1α and pHP1α. The most significant difference between the HP1α and pHP1α contact maps is the appearance of strong interactions between the NTEs and the hinge regions upon phosphorylation. These favorable contacts between the negatively charged phosphate groups and the positively charged hinge segment result in a more compact conformation of the HP1α dimer upon phosphorylation in our AA simulation.

To complement the AA simulations, which are computationally expensive and cannot be used to study the thermodynamic phase behavior,^41^ we next conducted coarse-grained (CG) simulations of HP1α and pHP1α dimers using the recently developed HPS-Urry and HPS-PTM models.^42,43^ In these simulations, the folded domains (CSD and CD) were kept rigid (to avoid protein unfolding) by applying a rigid body constraint^44^ while the rest of the chain remained flexible. This ensures that the CSD domains are kept fixed with respect to each other in a dimer configuration, while the folded domains are free to move (translate and rotate), and therefore are able to interact with other folded and disordered segments during the simulation. As shown in **Fig. 2d**, the R_g_ distributions sampled from our CG simulations are in very good agreement with the AA simulations. Moreover, compact configurations are populated for pHP1α with an average R_g_ = 2.86 ± 0.37 nm as compared to HP1α (R_g_ = 3.09 ± 0.53 nm). This compaction is mainly driven by the electrostatic attractions between the negatively charged NTE and the positively charged hinge regions as highlighted (magenta boxes) in the CG contact map (**Fig. 2e**). We also calculated the distance distributions between the two NTE segments of the homodimer HP1α (**Fig. S3b**), which provides insight into the collapsed and extended conformation probabilities. These distributions follow a similar pattern as observed for R_g_. The long tail of the distance distribution also suggests that unmodified HP1α can adopt much more extended conformations compared to its phosphorylated variant. The mean values of the NTE-NTE distance distributions from our simulations are similar to the D_max_ values^31^ obtained by published small angle X-ray scattering experimental data (5 – 6 nm) although the observed trends are reversed. The experimental data suggest that the NTEs in pHP1α are on average further apart compared to the NTEs in HP1α and that pHP1α exhibits a long tail of distance distributions. We, however, note that our simulations strictly reflect the behavior of one HP1α or pHP1α dimer, while the experimental concentrations (75 and 150 μM) may still be high enough to reflect oligomerization (see discussion of the oligomeric conformation of pHP1α below). In summary, both AA and CG models suggest that phosphorylation of the NTE leads to a more compact rather than an extended state of the homodimer mediated by electrostatic intra-dimer interactions between the NTE and hinge regions of the two monomers. The qualitative behavior from the CG model simulation is consistent with the AA simulation data, which provides further confidence in continuing with the CG model to study the phase behavior of HP1α proteins.

### The influence of phosphorylation on LLPS of HP1α

The observed compaction of pHP1α in AA and CG simulations is consistent with its enhanced LLPS propensity due to the known empirical correlations between single-molecule chain dimensions and phase separation propensity.^45,46^ Specifically, the favorable intramolecular contacts that dictate the compaction of a single chain are also important to establishing the inter-chain interactions that can stabilize the protein condensed phase. The advantage of the CG model is that it can be used to simulate the phase behavior and identify the molecular interactions within the condensed phase of HP1α and pHP1α. We conducted CG phase coexistence simulations of the homodimer using slab geometry^47,48^ as done in our previous work.^49^ As the coexistence density in the dilute phase (referred to as the saturation concentration C_sat_) is too low at 300K, we simulated the systems at 320K and plotted the protein density as a function of the z-coordinate that separates the dense phase from the dilute phase (**Figs. 2g-h**). We find that the NTE phosphorylation of HP1α lowers the saturation concentration significantly (by approximately sevenfold) as shown in the inset of **Fig. 2g**. In other words, pHP1α can undergo phase separation at a much lower protein concentration than HP1α, which is consistent with the experimental data and the relationship between protein collapse and phase separation propensity.

It is interesting to note that both the saturation concentration and the condensed phase concentration decrease upon phosphorylation (**Fig. 2g-h, Movies S3 and S4**). This observation suggests that there is an increase in the size of the condensate when the NTE is phosphorylated. We suspect that the increased net negative charge upon phosphorylation causes electrostatic repulsion within the condensed phase leading to expansion of the condensate. To test this hypothesis, we deleted 14 amino acids from the CTE of pHP1α and repeated the coexistence simulations. As the CTE carries significant negative charge, deletion of these residues reduces the overall negative charge of pHP1α without compromising the interactions of the phosphorylated NTE. The CG coexistence simulations for this construct show that the dense phase concentration significantly increased as expected (**Fig. S3c**). These observations are also in agreement with previous experimental studies where the deletion of the CTE lowers the saturation concentration for phase separation.^31^

To characterize the influence of phosphorylation on the interactions between HP1α dimers in the condensed phase, we calculated the intermolecular vdW contact maps of HP1α and pHP1α (**Fig. S4**). For the HP1α homodimer, the critical inter-dimer interactions relevant for phase separation are mainly driven by the electrostatic attraction between oppositely charged regions. Positively charged patches in the hinge from one dimer, KKYKK (beginning of the hinge), KRK (in the middle of the hinge), and KSKKKR (near the end of the hinge), act as hotspots to attract negatively charged regions from neighboring dimers, such as the EDEEE patch between the NTE and the CD, the EDAE patch in the CTE, and scattered acidic residues in the folded CD and CSD domains. (**Figs. 1 and S4a**). Upon phosphorylation, the considerable negative charge added to consecutive serine sites (S11-14) makes NTE-hinge region contacts more dominant in bridging neighboring pHP1α dimers (**Fig. S4b**). The result also shows a strong correlation between the interactions promoting LLPS of pHP1α and the interactions inducing the compaction of the pHP1α homodimer. While NTE-hinge contacts are enriched in LLPS of pHP1α, the contact propensity for other regions decreases. This is likely due to the increase in condensate size upon phosphorylation. Based on these observations, we hypothesized that phosphorylation promotes chain expansion in the condensed phase, allowing multivalent interactions through NTE-hinge bridging, but induces compact conformation of HP1α dimer in the dilute phase (as shown in the dimer AA and CG simulations in **Fig. 2**). To test this, we calculated R_g_ and NTE-NTE distributions of the homodimer in HP1α and pHP1α CG coexistence simulations. We found that pHP1α adopts a more extended conformation in the condensed phase, with R_g_ = 4.24 ± 0.81 nm, compared to R_g_ = 4.03 ± 0.77 nm of HP1α (**Fig. 2i**). The NTE-NTE distance distributions also follow a similar behavior as observed for Rg (**Fig. S3d**). These results agree with the published small angle X-ray scattering (SAXS) data that indicate a more elongated conformation of pHP1α.^31^

### The effect of peptide ligands on LLPS of pHP1α

Electrostatic interactions that involve the phosphorylated NTE of one dimer and the hinge region of another dimer appear to be the driving force behind LLPS of pHP1α. HP1α, however, can also simultaneously interact with multiple binding partners through the CSD dimer interface.^24^ These interactions may change the electrostatic landscape of HP1α and thus lead to modulation of LLPS. Larson et al.,^31^ for example, showed that peptides from the binding partners shugoshin (Sgo1) and lamin B receptor (LBR) can affect the saturation concentration of pHP1α droplet formation. This was attributed to changes in the overall interaction patterns of the phosphorylated NTE, hinge, and CTE regions. Using our combined experimental and computational approach, here we set out to decouple the contributions of specific and non-specific ligand interactions and to provide a comprehensive picture of how peptide ligands may affect LLPS of pHP1α.

For this purpose, we chose a set of four peptides from three known binding partners of HP1α, namely Sgo1, LBR, and chromatin assembly factor 1 (CAF1).^50–52^ We also selected a peptide corresponding to the αN helix of H3 which contains a PXVXL-like motif and can interact with HP1α in the non-nucleosome context.^53,54^ All peptides can bind the CSD-CSD homodimer as confirmed by NMR titration experiments (**Fig. S5-S9**). As previously reported, the symmetric HP1α-CSD homodimer produces well-resolved ^1^H-^15^N HSQC spectra where each cross-peak reflects an amide ^1^H-^15^N pair in the protein (**Fig. S5**).^53,55^ Upon addition of the peptides, CSD residues 164-174, which contain the peptide binding interface, either disappeared or shifted in the spectrum (**Fig. S6-S9**), consistent with previous observations.^53,56^ Several additional peaks were also perturbed in each spectrum, indicating some allosteric structural effects upon peptide binding. Based on the cross-peak splitting profiles, the symmetry of the homodimer seemed to be broken once the peptides were bound to the homodimer binding interface. NMR titration experiments were also performed with the slightly longer CSD-CTE homodimer, with similar results (**Fig. S6-S9**). Published binding studies indicate that Sgo1 and CAF1 display tighter binding (K_d_ of ∼ 0.2 – 0.4 μM), while LBR and H3 have a Kd of ∼ 3 μM and 58 μM, respectively (**Table S2)**.^31,53^ Overall, our results agree well with previous literature and the binding trends appear to be very similar between dimer constructs that contain the CSD domains alone, the CSD-CTE, or full-length HP1α.

We then investigated the effect of peptide addition on LLPS by CG co-existence simulations (**Fig. 3b-c, Fig. S10**). We used pHP1α homodimers and peptides at a 1:1 ratio and calculated the condensed and saturation concentrations of the protein. These simulations were initiated from a high-density slab, where HP1α homodimers and peptides were mixed and able to interact with each other via the same CG HPS-Urry model that was used for interactions between the HP1α proteins (**Fig. 3b, Movie S5**). We found that the Sgo1 and H3 peptides lowered the saturation concentration of LLPS (**Fig. 3c**) and were mostly found in the dense phase. In contrast, the addition of LBR and CAF1 peptides inhibited phase separation. These peptides interacted weakly with HP1α, and hence were abundant in the dilute phase (**Fig. 3b-c**).

**Figure 3.**
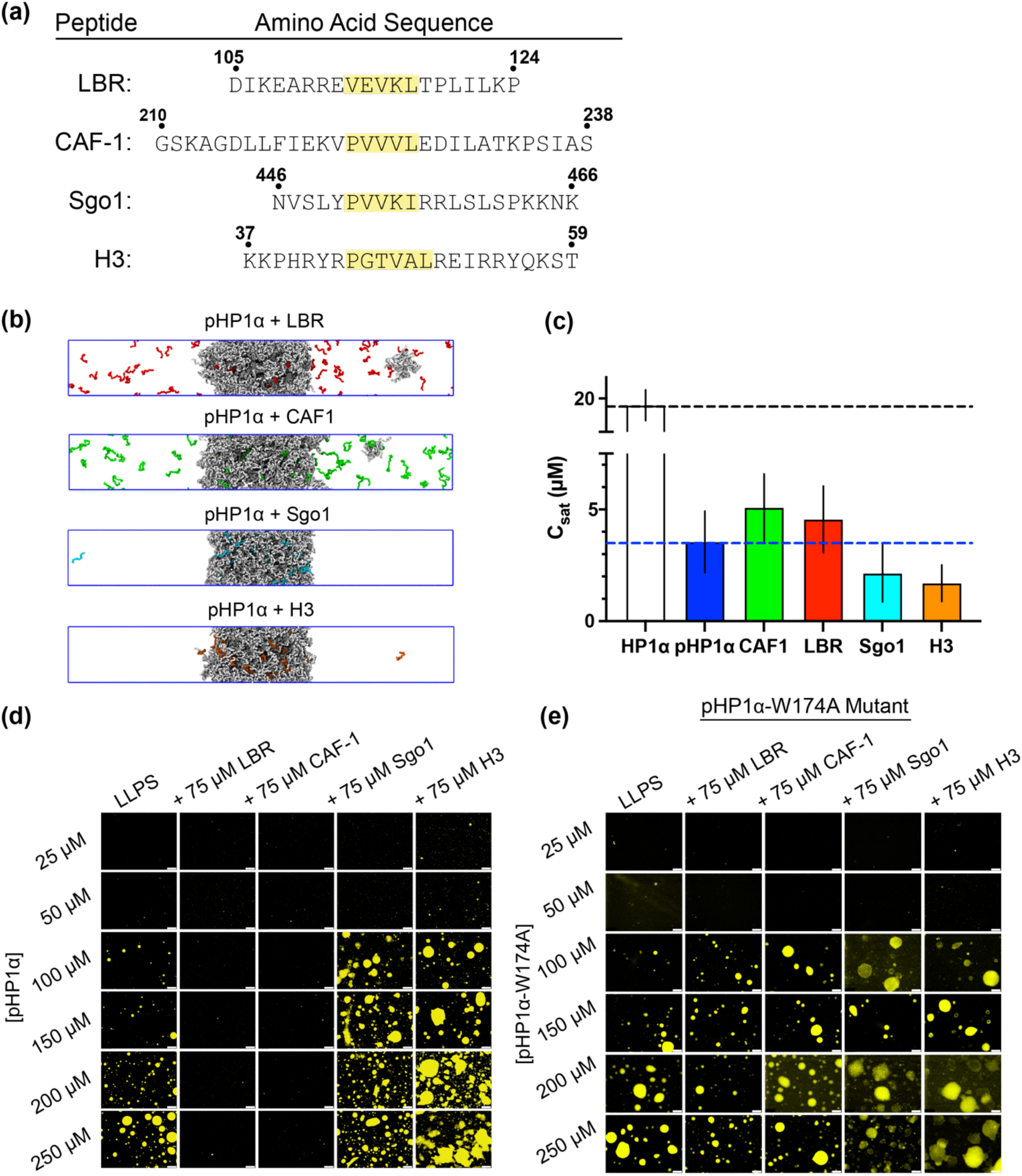
Influence of peptide ligands on LLPS of pHP1α. **(a)** Amino acid sequence of the peptide ligands used in this study (LBR, CAF-1, Sgo1, and H3). Their PXVXL or PXVXL-like motifs are highlighted in yellow. **(b)** Snapshots of slab simulations of pHP1α and peptides at 1:1 ratio of peptide to pHP1α. **(c)** Saturation concentration measured in the slab simulations at 320K. The first two cases, HP1α (white) and pHP1α (blue), were simulated without the addition of peptides; the rest of the simulations were performed in the presence of peptides. The error bars represent the standard deviation from triplicate sets of simulations. **(d)** and **(e)** Fluorescence microscopy images of LLPS of pHP1α and pHP1α-W174A with and without 75 μM peptide, as indicated on top of each column. Fluorescent droplets were visualized with 0.6 μM Cy3-labeled pHP1α added to the LLPS samples. The white scale bar represents 100 μm.

We next proceeded with evaluating the effect of the peptides on pHP1α phase separation using microscopy. To visualize LLPS, we used 0.6 μM of pHP1α labeled with the fluorescent dye Cy3 on an engineered cysteine residue at the C-terminus (pHP1α-Cy3). We incubated varying concentrations of pHP1α with 75 μM peptide and imaged the samples by Cy3 fluorescence and by brightfield microscopy (**Fig. 3d** and **Fig. S11**). Consistent with the simulation results, the H3 and Sgo1 samples appeared to form droplets more efficiently than pHP1α alone, while the LBR and CAF1 peptides perturbed pHP1α phase separation for all tested pHP1α concentrations.

Based on binding affinity, the peptides that we studied can be divided into slightly tighter binders (Sgo1 and CAF1) and slightly weaker binders (LBR and H3).^31,53^ However, both the computational and experimental results indicate that their effect on LLPS is based on charge rather than binding affinity. Sgo1 and H3 are rich in lysine and arginine residues and carry a high net positive charge of +5.9 and +7, respectively, while LBR and CAF1 have a weak net charge of +0.9 and -1.1, respectively. Taken together, our results suggest that positively charged peptide ligands such as Sgo1 and H3 enhance LLPS, while ligands with a weak net charge such as LBR and CAF1 destabilize the condensed phase.

### The contribution of specific peptide-pHP1α interactions in LLPS

While peptide ligands appear to modulate LLPS based on their net charge, it is not clear if modulation is due to specific binding to the CSD-CSD dimer or to non-specific electrostatic interactions with other segments of the pHP1α dimer. To dissect the contributions of specific and non-specific interactions, we prepared pHP1α with the W174A mutation (pHP1α-W174A).^53^ This construct eliminates a key tryptophan residue that is important in the recognition of the PXVXL or PXVXL-like motifs in the binding partners. NMR experiments confirmed that this mutation does not perturb the overall structure of the CSD-CTE homodimer, while gel shift assays verified that peptide binding through the dimerization interface has been abolished (**Fig. S12**).

Using pHP1α-W174A, we repeated the fluorescence microscopy experiments described above to assess the effect of peptides on LLPS in a context where specific peptide-pHP1α interactions were eliminated. These experiments indicated that the peptides no longer had a discernible effect on LLPS (**Fig. 3e** and **Fig. S13**). While microscopy provides a qualitative picture of the effects of each peptide on pHP1α phase separation, we also sought out to obtain a more quantitative view of the observed trends. LLPS was induced in the presence of each peptide and the sample was centrifuged in order to separate the droplet and the supernatant phase. We then measured the A280 absorbance of the supernatant solution. Since all peptides used in our study have low extinction coefficients at 280 nm (**Table S3**), the A280 absorbance of the supernatant was attributed to pHP1α and was used to determine the concentration of the protein in that phase. This provided a quantitative measure of the propensity of pHP1α to undergo LLPS in the presence of different peptides. First, we kept the concentration of peptide constant while increasing the concentration of pHP1α (**Fig. 4a**). Similar to the microscopy experiments, Sgo1 and H3 appeared to enhance LLPS while LBR and CAF-1 behaved similarly to the control samples that do not undergo LLPS (pHP1α at 300 mM KCl). In the absence of binding, i.e. when pHP1α-W174A was used, the Sgo1, LBR and CAF1 peptides did not affect LLPS, while the H3 peptide still enhanced LLPS (**Fig. 4b**). We note that in these experiments the pHP1α-W174A construct appears to phase separate better compared to pHP1α. This may be due to enhanced propensity for LLPS due to the mutation or due to higher levels of phosphorylation which is hard to control when performed enzymatically.

**Figure 4.**
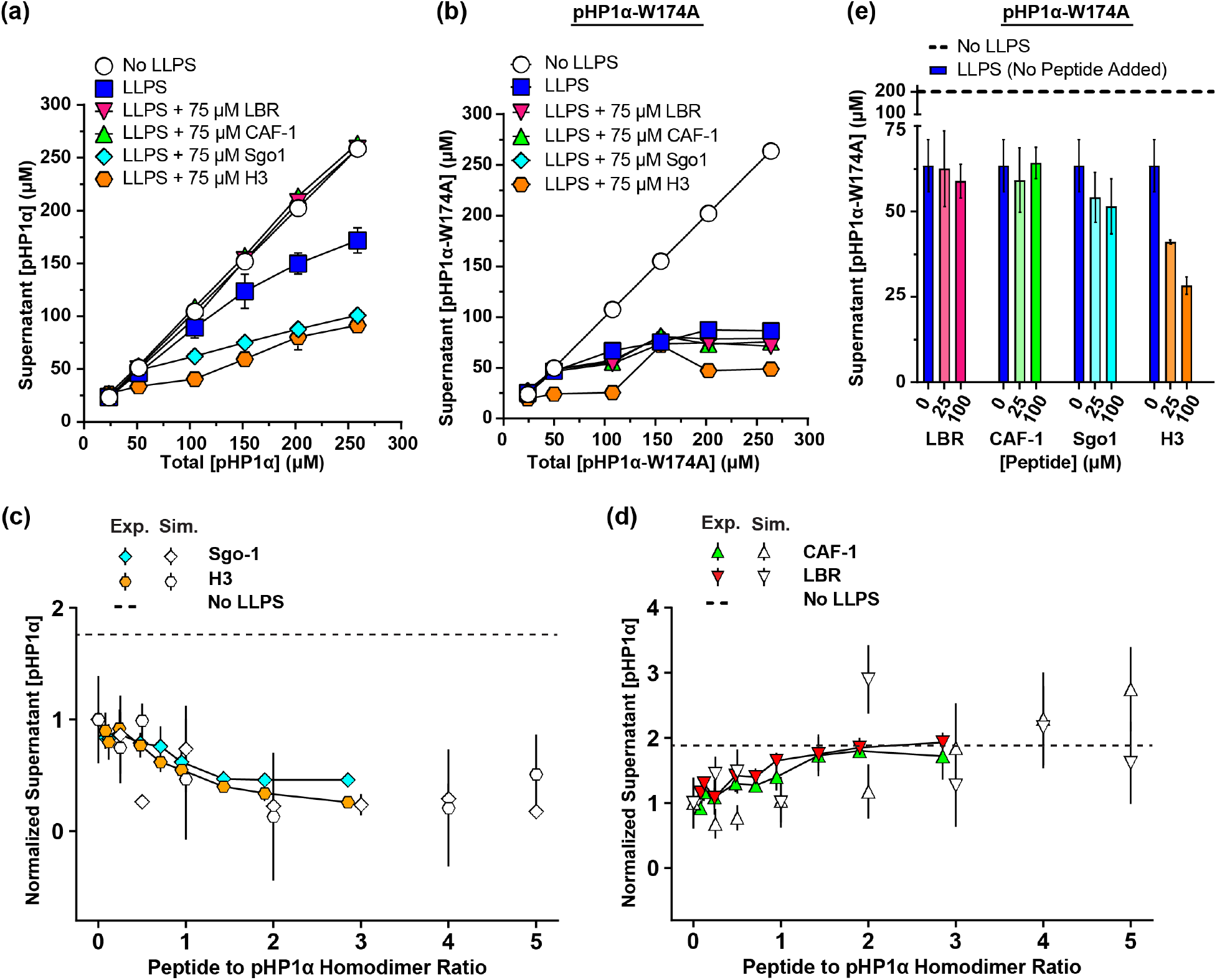
LLPS of pHP1α and pHP1α-W174A in the presence of peptide ligands. **(a)** Concentration of pHP1α in the supernatant after LLPS using a range of starting pHP1α concentrations and 75 μM peptide. **(b)** Concentration of pHP1α-W174A in the supernatant after LLPS using a range of starting pHP1α-W174A concentrations and 75 μM peptide. **(c, d)** Normalized supernatant pHP1α concentration of LLPS with and without peptide. Experimental data are shown with color symbols, while the simulated data are depicted with white symbols. The dashed lines represent the experimental ratio that will be calculated if LLPS does not occur. Each data point is normalized to the supernatant concentration of pHP1α in the absence of peptide. **(e)** pHP1α-W174A concentration in the supernatant after LLPS using 200 μM pHP1α-W174A with 25 and 100 μM peptide, respectively. The dashed line represents the experimental concentration that will be measured if LLPS does not occur. The error bars of the experimental data in **(a-e)** and the computational data in **(c, d)** represent the standard deviation from triplicate sets of experiments and simulations, respectively.

Since the peptides have different binding affinities to the homodimer interface of pHP1α, the concentration of unbound peptides might also influence LLPS through non-specific mechanisms such as interactions with the hinge, NTE or CTE. Therefore, we repeated the absorbance experiments by keeping the concentration of pHP1α constant while using different ratios of peptide to pHP1α homodimer. We expected that non-specific mechanisms would manifest at higher ratios of peptide to pHP1α homodimer and would perturb the concentration of pHP1α in the supernatant. For the positively charged Sgo1 and H3 peptides, the concentration of pHP1α in the supernatant decreased at low peptide to dimer ratios, and then remained constant as more peptide was added (**Fig. 4c**). CAF-1 and LBR on the other hand, promoted the increase of pHP1α in the supernatant until LLPS was completely abolished (**Fig. 4d**). These trends were also confirmed quantitatively by our CG co-existence simulations performed at increasing ratios of peptide to pHP1α (**Fig. 4c,d**), which is remarkable given that no attempt was made to modify the previously proposed model to match the experiment and no fitting is involved. Interestingly, we also observed a weak non-monotonic trend in the saturation concentration which appears to increase at high concentrations of positively charged peptides (**Fig. 4c**). This may be due to electrostatic repulsion when the addition of peptides exceeds the amount required to neutralize the net charge of pHP1α. We have also observed this trend experimentally in different batches of pHP1α (**Fig. S14**).

In the absence of specific binding (i.e. when pHP1α-W174A was used), positively charged peptides can still promote LLPS at sufficiently high peptide concentrations (**Fig. 4e**). This is particularly evident for H3. We also performed experiments with a highly positively charged peptide from the histone H4 tail which does not contain a PXVXL-like motif and is not known to bind the CSD dimer interface. This peptide affected LLPS in a manner similar to the H3 peptide (**Fig. S15**). Therefore, it appears that the charge of the peptide is an important driver in LLPS. Highly positively charged peptides such as H3 and H4 can promote pHP1α phase separation even in the absence of specific interactions with the CSD dimer interface. This is also supported by the CG co-existence simulations where no specific interactions were encoded between the peptide and the protein dimer. In this case, the peptides can accumulate within the droplet through attractive non-specific electrostatic interactions to help balance the highly negative charge of the condensed phase. The specific binding interaction, however, is important for the action of peptides with lower charge as a means to increase their local concentration within the droplet.

### Computational studies of HP1α LLPS in the presence of DNA

Experimental studies have shown that HP1α can interact with DNA through patches of basic residues in the hinge, which allows HP1α dimers to bridge different regions of DNA and to induce DNA compaction.^27,31^ HP1α-DNA interactions also initiate an increase in the local concentration of HP1α and possibly promote HP1α-HP1α interactions to form higher order oligomers leading to condensate formation. DNA-driven phase separation of HP1α is mainly governed by electrostatic interactions, as raising the level of monovalent salts increases the saturation concentration.^27^ In addition, HP1α-DNA condensation also depends on DNA length, and the concentration of nucleic acid and HP1α. It has been reported that both HP1α and pHP1α can undergo LLPS in the presence of DNA.^27,32^

There are ongoing efforts to develop and evaluate a nucleic acid (DNA/RNA) CG model to study protein-nucleic acid interactions and the role DNA plays in LLPS.^57–60^ Here, we use a model that separates the nucleotide into two beads; one bead represents the sugar-phosphate backbone, carrying an overall -1 charge, and the other represents the base but differentiates between bases ADE, THY, CYT, and GUA in their respective stacking and hydrogen bonding interactions. Using our CG nucleic acid model, we studied the effects of DNA addition on the LLPS of HP1α by conducting CG coexistence simulations of HP1α homodimers containing a small mole fraction of 205 bp dsDNA, 0.02 and 0.038, at 320 K. This system can be reasonably compared to an experimental system containing 50µM/100 µM of HP1α with 1.5µM of the same 205 bp dsDNA. In our simulations, dsDNA partitions into the droplet and promotes LLPS in a concentration dependent manner (**Fig. 5a, Movie S6**). We find that the addition of dsDNA at a mole fraction of 0.038 lowers the saturation concentration nearly ten-fold as compared to HP1α alone (**Fig. 5a inset**). We also conducted simulations of HP1α homodimers containing 0.074 mole fraction of dsDNA, however, a tremendous increase in the overall negative charge of the system weakened the condensed phase network of HP1α and resulted in the formation of void volume within the simulated slab (**Movie S6**).

**Figure 5.**
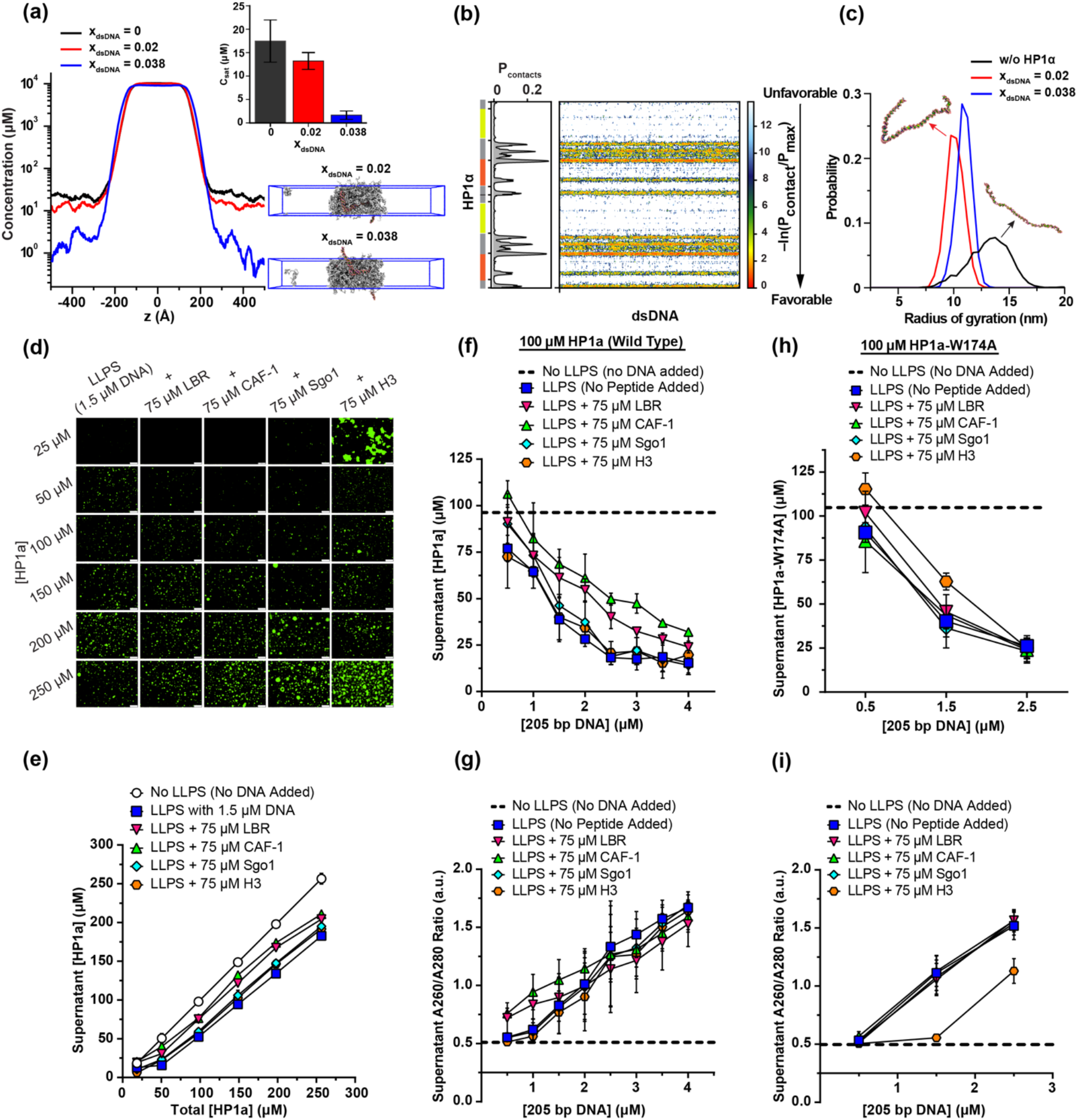
LLPS of HP1α with DNA. **(a)** Density profiles, saturation concentrations (inset) and snapshots of HP1α condensates as a function of the added DNA fraction, as determined in the CG coexistence simulations. **(b)** Intermolecular contacts between HP1α and DNA within the condensed phase. Preferential interactions between HP1α and DNA are shown in red. **(c)** Radius of gyration distribution of DNA within HP1α condensates. **(d)** Fluorescence microscopy images of LLPS of HP1α using 1.5 μM of 205 bp DNA, with and without 75 μM peptide as indicated on top of each column. Imaging was performed with 0.1 μM YOYO-1 dye which binds DNA. The white scale bar at the bottom right of each image represents 50 μm. **(e)** HP1α concentration in the supernatant after LLPS using a range of starting HP1α concentrations and 1.5 μM of DNA with and without 75 μM of peptide. **(f)** HP1α concentration in the supernatant after LLPS using 100 μM of HP1α and a range of DNA concentrations, with and without 75 μM peptide. **(g)** The A260/A280 ratio of the supernatant after LLPS using 100 μM of HP1α and a range of DNA concentrations, with and without 75 μM peptide. **(h)** HP1α-W174A concentration in the supernatant after LLPS using 100 μM of HP1α-W174A and a range of DNA concentrations, with and without 75 μM peptide. **(i)** The A260/A280 ratio of the supernatant after LLPS using 100 μM of HP1α-W174A and a range of DNA concentrations, with and without 75 μM peptide.

To characterize the protein-DNA molecular interactions responsible for promoting LLPS in our system, we computed the inter-molecular contact map based on vdW contacts formed between HP1α and dsDNA as a function of residue number (**Fig. 5b**). The contact map agrees with experimental observations^27^ where prominent hinge-DNA interactions are mediated by the KKYKK (beginning of the hinge), KRK (in the middle of the hinge), and KKK (near the end of the hinge) patches. In addition, we find two more contact-prone regions, one in the disordered NTE due to patch KKTKR, and one in the CD due to the patch RRVVK. These observations confirm that electrostatic interactions between the negatively charged sugar-phosphate backbone of the DNA and the positively charged lysine/arginine-rich hinge and NTE disordered regions of HP1α are the driving force behind LLPS of HP1α in presence of DNA.

To gain insight into the effect of HP1α -DNA interactions on the DNA size, we also computed the distribution of the radius of gyration (Rg) of the dsDNA chains in the presence and absence of HP1α homodimers (**Fig. 5c**). This analysis shows that dsDNA adopts a more collapsed state when the DNA chains are interacting with HP1α, suggesting that dsDNA gets compact while aiding LLPS of HP1α. This observation agrees with previous experimental observations based on DNA curtain compaction assays.^27,31^ Overall, our CG model of DNA-driven HP1α LLPS reproduces experimental trends and implicates hinge-DNA interactions as a main driver of phase separation.

### The effect of peptides on DNA-driven LLPS of HP1α

Considering the electrostatic nature of the involved interactions, we wondered how peptide ligands would affect the phase separation of non-phosphorylated HP1α with DNA. For this purpose, we induced LLPS in the presence of the same 1.5 μM 205 bp DNA segment described above, with and without 75 μM peptide. Qualitative microscopy images did not reveal substantial differences between the control and samples containing LBR, CAF-1 or Sgo1 (**Fig. 5d** and **Fig. S16**). On the other hand, H3 initially formed large clumps that evolved into liquid droplets as the concentration of HP1α was increased. As H3 is highly positively charged, it is likely that its interactions with DNA dominate at low HP1α concentrations. Upon addition of HP1α, H3 can interact with both DNA and the dimer interface, and as the HP1α-DNA interactions become more prevalent, LLPS can occur. Similar results were observed with the H4 peptide (**Fig. S17 and S18**). We complemented these experiments with quantitative A280 measurements of the concentration of HP1α in the supernatant after LLPS with DNA (**Fig. 5e**). These experiments suggest that LBR and CAF-1 have a slight tendency to perturb LLPS, while Sgo1 and H3 have no effect.

We repeated these experiments holding the concentration of HP1α and peptide constant, while varying the amount of DNA (**Fig. 5f**). The trends here were the same, with LBR and CAF-1 consistently leading to a higher concentration of HP1α in the supernatant, while Sgo1 and H3 had no effect. The A260/A280 ratio can be used as a qualitative measure of the presence of DNA in the supernatant (**Fig. 5g**). Measurements of this ratio indicate that there were no significant differences in the amount of DNA that was present in the supernatant when different peptides were added. This suggests that the observed impact of LBR and CAF-1 on LLPS is not due to lower concentrations of DNA in the droplet phase. To further explore the ability of the HP1α dimer-peptide complex to influence LLPS, we used the W174A mutant and induced LLPS in the presence of 75 μM peptide, 100 μM HP1α-W174A, and varying concentrations of DNA (**Fig. 5h**). In this context, LBR and CAF-1 had no significant impact on the concentration of HP1α-W174A in the supernatant. This implies that their specific binding interaction with HP1α is responsible for LLPS modulation.

The only peptide that affected LLPS of HP1α-W174A was H3 (**Fig. 5h**). This peptide also appeared to sequester DNA from the supernatant as suggested by the lower A260/A280 ratio (**Fig. 5i**). Similar behavior was observed with the H4 peptide as well (**Fig. S19**). These highly positively charged peptides appear to compete with the hinge region of HP1α for access to DNA, especially at low DNA concentrations. When H3 binds to the homodimer interface through its PXVXL motif, the effective concentration of peptide available for DNA interactions will be reduced and H3 should not significantly perturb DNA-driven HP1α LLPS. When this interaction is abolished, as is the case for the HP1α-W174A sample, H3 can compete with HP1α for DNA sites more efficiently. In summary, our peptide studies of HP1α and DNA phase separation reveal a complex picture where LLPS can be modulated by a careful balance between peptide-HP1α specific binding interactions and competition between peptide and HP1α for non-specific interactions with DNA.

## Discussion

HP1α is an essential component of heterochromatin domains that contain repetitive DNA sequences, have distinct replication timing, and exhibit low levels of transcription.^3,61^ While these domains remain stable over time, the HP1α proteins inside are characterized by high mobility and fast exchange times when interacting with chromatin.^27,62,63^ It has been hypothesized that phase separation of HP1α is essential in the formation and maintenance of these domains.^5,31,64^ *In vitro* experiments have shown that this process can be modulated by several factors: 1) phosphorylation of the serine patch located in the NTE region of the protein; 2) multivalent interactions with DNA and chromatin; and 3) through ligands that target a specific binding site located at the CSD-CSD dimer interface.^27,31,32^ These factors can modulate the phase separation of HP1α in a positive or negative manner and may interact together in complex ways to finetune the biophysical properties of heterochromatin environments. Here, we provide a systematic investigation of the effect of these modulators on HP1α phase separation *in vitro* and describe the complex molecular interaction landscape that controls the formation of HP1α condensates.

At physiological pH, wild-type HP1α is overall a negatively charged protein, with negative charges concentrated in the CD, CSD, and CTE domains (**Fig. 1a**). Positively charged arginine and lysine residues are clustered in the disordered hinge region while the NTE has a small positive charge. As our atomistic simulations of the HP1α homodimer suggest, in this context, the predominant contacts are between the NTE regions, the NTE and the CD domain, and between the CTE and hinge (**Fig. 2c**). Upon phosphorylation, the charge on the NTE changes dramatically, and NTE-hinge region contacts become much more prevalent. CTE-hinge region contacts are still observed, implying that the negatively charged CTE and NTE regions may compete for access to the positively charged hinge region. Under dilute conditions, such as those represented by the atomistic and coarse-grained simulations, these contacts lead to a more compact state of pHP1α compared to the unmodified protein (**Fig. 2b** and **Fig. 6a**). Under crowding conditions, however, intradimer NTE-hinge contacts can be efficiently replaced by interdimer interactions, leading to a more extended pHP1α conformation (**Fig. 2i, S4c** and **Fig. 6a**). Our observations are consistent with the model of pHP1α LLPS proposed by Larson *et al*.,^31^ although we note that the effect of NTE phosphorylation on the overall conformation of HP1α may depend on protein concentration.

**Figure 6.**
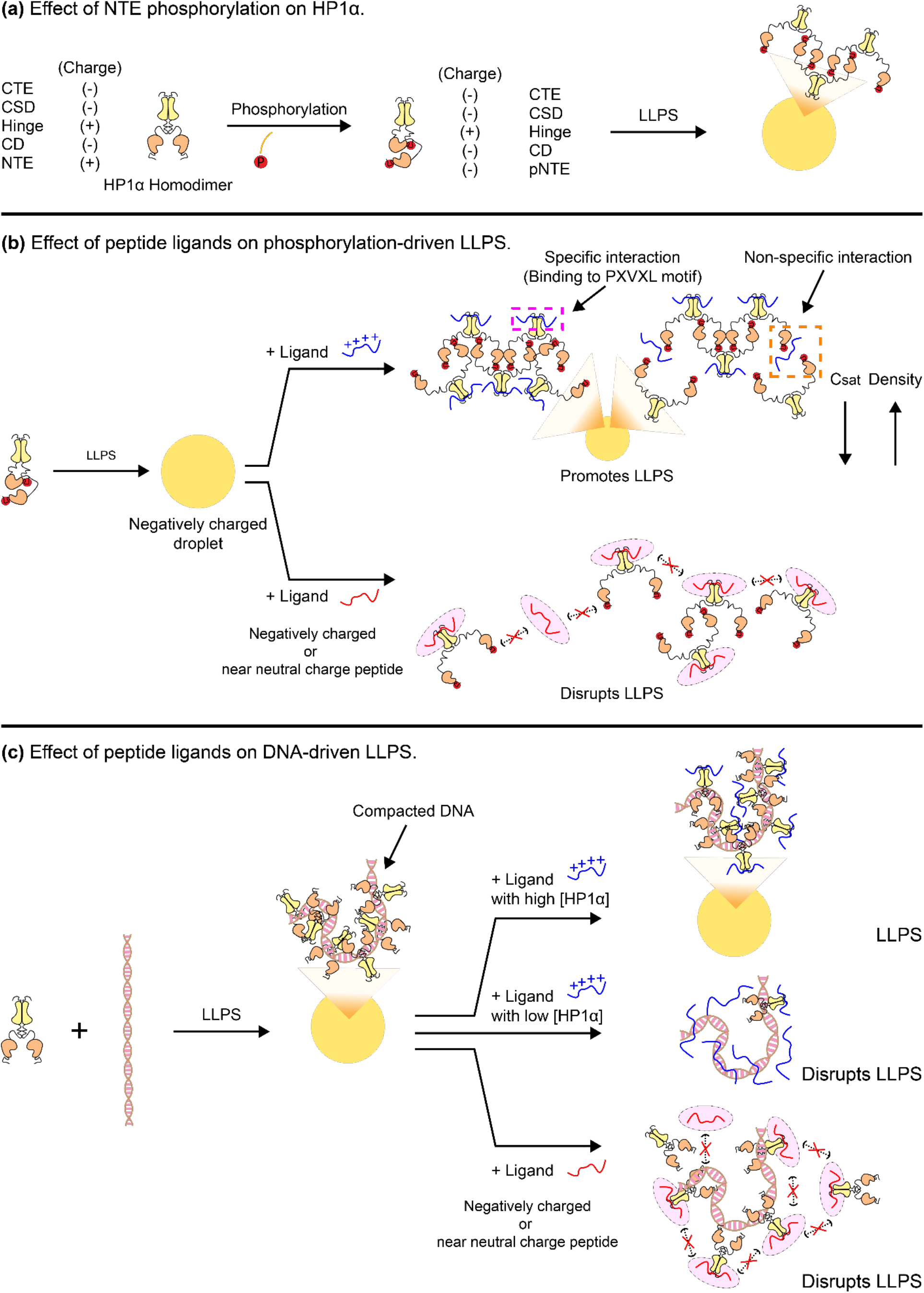
A complex network of specific and non-specific interactions controls HP1α LLPS. **(a)** NTE phosphorylation leads to a compact HP1α conformation under dilute conditions and an extended conformation during LLPS. In both cases, interactions are driven by the phosphorylated NTE and the hinge regions of the protein. **(b)** The effect of peptides during phosphorylation-driven LLPS of HP1α. Positively charged peptides can promote pHP1α LLPS by neutralizing the negative charge of the droplets (and increasing their density), by non-specific interactions that cross-link pHP1α dimers, and by specific interactions that redistribute the charge patterns on pHP1α. Peptides with low charge, on the other hand, disrupt pHP1α LLPS by adding negative charge to the pHP1α dimer and/or by screening phosphorylated NTE-hinge interactions. **(c)** The effect of peptides during DNA-driven LLPS of HP1α. Positively charged peptides may disrupt LLPS at low HP1α concentrations by competing for binding sites with DNA. Negatively charged or neutral peptides can disrupt LLPS by screening hinge-DNA interactions.

The observation that NTE phosphorylation enhances LLPS of HP1α is somewhat counterintuitive as this modification adds a more negative charge to an already negatively charged system. However, the addition of four adjacent phosphate groups at the NTE replaces the slightly positive charge of this region with a highly negatively charged patch, while the hinge region becomes the only segment that carries positive charge. Theoretical models have predicted that changes in charge patterning can have profound effects on the driving forces of phase separation.^65^ In particular, a sequence that contains blocks of charged residues has a higher tendency to undergo LLPS compared to a sequence with the same overall charge but with alternating positively and negatively charged residues. In essence, phosphorylation creates a block of strong negative charge on the N-terminus and a block of strong positive charge on the hinge region, ensuring stronger electrostatic attractions between the two regions in the extended interdimer network. This charge patterning then leads to the lower saturation concentration for pHP1α as observed experimentally and computationally. However, phosphorylation significantly decreases the overall charge of pHP1α and the resulting condensed phases have a large negative charge. Therefore, the favorable NTE-hinge region interactions are counterbalanced by repulsive interactions between the other regions of the protein. This results in a larger radius of gyration and lower density of the condensed phase as compared to unmodified pHP1α. It should also be noted that the strict interpretation of the charged patterning model as it relates to pHP1α is complicated by the presence of folded domains that may contribute to LLPS through other types of interactions. For example, in some crystal structures, CSD-CSD dimers can interact with each other through a hydrophobic β-sheet interface.^19^ While such interactions are likely rare under dilute conditions, they may become more prevalent at high protein concentrations and may compete with the attractive and repulsive electrostatic forces promoted by phosphorylation.

Overall, the molecular effects of NTE phosphorylation on HP1α LLPS appear complex. Phosphorylation redistributes the charge patterns in the protein sequence and strengthens the interactions of the NTE and the hinge regions. At the same time, it provides an additional negative charge that introduces repulsion among other regions of the protein and leads to an extended protein conformation at high concentrations (**Fig. 6a**). These competing effects can be modulated by HP1α ligands that interact specifically with the symmetric CSD-CSD dimer interface. Our studies indicate that highly positively charged peptide ligands promote phase separation of pHP1α while peptides with neutral or slightly negative charge are disruptive to LLPS (**Fig. 3**). In addition, we find that highly positively charged peptides such as H3 and H4 can enhance LLPS even in the absence of specific binding, while peptides with low charge require binding to the CSD-CSD dimer interface to modulate phase separation at the tested concentrations.

The observed effects of the peptide ligands are consistent with several non-exclusive mechanisms (**Fig. 6b**). First, positive charge may be required to stabilize the highly negatively charged pHP1α droplets. Assuming only non-specific interactions, e.g. when the pHP1α-W174A mutant is used, this mechanism would predict that only positively charged peptides would modulate LLPS while peptides with negative charge would be incompatible with droplet entry. However, when specific interactions are possible, the accumulation of additional negative charge on the pHP1α dimer would make the droplets even more negative, leading to their eventual dissolution. We also note that at sufficiently high concentrations of positively charged ligands, the relationship should be reversed and LLPS would be disrupted, as demonstrated both by our experimental and computational studies (**Fig. 4c** and **Fig. S14a**). Second, free peptides in the droplet may affect the crosslinking interactions between pHP1α homodimers. For example, in our coarse-grained simulations, positively charged peptides have a strong preference for the phosphorylated NTE (**Fig. S10**) and thus may serve as an additional motif that provides the opportunity for more multivalent interactions. Negatively charged or neutral peptides, on the other hand, may screen the NTE-hinge interactions that drive LLPS. This mechanism of LLPS modulation would require sufficient amounts of free peptide in the droplets, e.g. at high peptide to pHP1α homodimer ratios for stronger specific ligands such as LBR, CAF-1 and Sgo1, or for the H3 and H4 peptides used in our study which bind to the specific dimer interface weakly or not at all (**Fig. 4** and **Fig. S15**). Finally, the specific binding of the peptide PXVXL motif to the dimer interface can alter the dynamics and interaction patterns of the CTE region. For example, Larson *et al*., hypothesized that positively charged peptides like Sgo1 can disrupt CTE-hinge interactions, resulting in less competition with the NTE.^31^ Alternatively, a negatively charged peptide ligand at the dimer interface may favor CTE-hinge interactions that disrupt LLPS. In this case, abrogating specific binding through mutation should eliminate the peptide effects on LLPS as observed for CAF-1, LBR and Sgo1. These potential mechanisms illustrate the rich network of interactions that can be exploited by ligands in finetuning the biophysical and functional properties of pHP1α.

Similar to phosphorylation driven LLPS, wild type HP1α also relies on its hinge region to undergo LLPS with DNA. Previous literature has suggested that in phase-separated HP1α-DNA condensates, HP1α remains highly dynamic while the DNA polymers are compacted and constrained through multivalent interactions with the HP1a hinge region.^27^ Our coarse-grained simulations agree well with this model and capture both the hinge-DNA contacts and the compacted state of DNA in the condensed phase (**Fig. 5)**. This two-component condensate system responds to peptide ligands in different ways depending on 1) the propensity of the peptide to interact specifically with the CSD-CSD homodimer interface, 2) the peptide affinity for DNA, and 3) the concentration of HP1α homodimers and DNA. The effects of positively charged peptides such as Sgo1, H3 and H4 range depending on the strength of the specific interaction with the CSD-CSD homodimer interface (**Table S2**). As the highest affinity ligand investigated in our study, the Sgo1 peptide is likely sequestered at the homodimer interface and does not significantly compete for binding with DNA. The H3 peptide has no effect on LLPS when DNA and HP1α are abundant, while H4 sequesters DNA to form amorphous aggregates that disrupt LLPS. Negatively charged or neutral peptides (e.g. CAF-1 and LBR), on the other hand, may provide some screening of the hinge region leading to slight disruption of LLPS. This effect is also most likely K_d_ dependent. Therefore, it appears that DNA-driven LLPS of HP1α can be disrupted by both negatively and positively charged ligands and its regulation requires a careful balance of the ligand affinity towards HP1α or DNA (**Fig. 6c**).

Our studies provide a molecular window into the interactions that drive and modulate HP1α phase separation *in vitro* and *in silico*, where we could dissect the effect of modulators and ligands one at a time. Even in this simplified context, a complex picture emerges where hinge-NTE and hinge-DNA crosslinking interactions must be carefully balanced by post-translational modifications and/or HP1α binding partners. The differential effects of peptide ligands on phosphorylation-driven and DNA-driven LLPS of HP1α may also have important biological consequences. Phosphorylation-driven condensation can be modulated both in the positive and negative direction, while DNA-driven phase separation can be disrupted by negatively charged ligands and competition for binding sites on the DNA. In cells, LLPS would also be modulated by other components such as the HP1β and HP1γ paralogs that can dissolve HP1α droplets,^27^ as well as the properties of chromatin polymers and post-translational modifications on the histone proteins.^66,67^ Nonspecific DNA-hinge contacts, for example, will be complemented by specific CD-H3 K9me2/3 interactions that increase the residence time of HP1α near chromatin.^26,62,63,68^ This increased multivalency may provide buffering capacity against disruptions by HP1α interaction partners and other heterochromatin components. Interestingly, pHP1α has a higher affinity for H3 K9me3, while displaying a lower affinity for DNA compared to the wild-type protein.^33,69^ pHP1α can also undergo LLPS with DNA and nucleosome arrays at similar concentrations as HP1α.^66^ Future experiments will no doubt reveal the molecular basis and functional consequences of these observations. More work is also needed in understanding the cellular function of NTE-phosphorylation, including the players that are involved in its regulation and its consequences on gene silencing and activation.

## Methods

### All-atom MD simulation protocol and analysis

We used the Amber99SBws-STQ force field with improved residue-specific dihedral correction,^34^ TIP4P/2005 water model^35^ and improved salt parameters from Lou and Roux^70^ for all systems. Force field parameters for phosphorylated serine were obtained from previous studies.^71^ Phosphorylated serine residues (S11-14) were modeled using Charmm-GUI.^72^ Energy minimization and equilibration were performed using GROMACS 2020.^73^ Each of the dimer systems (HP1α and pHP1α) was placed into an octahedral box of 15 nm length. The energy was first relaxed in vacuum, then the system was solvated with TIP4P/2005 water molecules and the energy was minimized further. The steepest descent algorithm was used in both energy minimization steps. To mimic the salt-concentration used in the experiment (0.075 M), Na^+^ and Cl^−^ ions were added along with additional Na^+^ counter ions to achieve electrical neutrality. NVT (canonical ensemble) equilibration was performed using the Nose-Hoover thermostat^74^ with a coupling constant of 1.0 ps. to stabilize the system temperature at 300 K. NPT (isothermal-isobaric ensemble) equilibration was performed using the Berendsen barostat^75^ with isotropic coupling with a constant of 5.0 ps to achieve a system pressure of 1 bar. All production simulations were performed using OpenMM 7.6^76^ in the canonical ensemble at 300 K using the Langevin middle integrator^77^ with a friction coefficient of 1 ps^−1^. Masses of all hydrogen atoms were increased to 1.5 amu which allowed for a simulation timestep of 4 fs. Constraints were applied to all hydrogen-containing bonds using the SHAKE algorithm.^78^ Short-range non-bonded interactions were calculated based on a cutoff radius of 0.9 nm. Long-range electrostatic interactions were treated using the PME method.^79^ Errors for Rg were estimated using block averages with five blocks. We calculated the contact map using the method described in the previous study.^80^ A vdW contact between two residues was considered formed if at least one atom from one residue was within 6Å distance from an atom in the other residue.

### CG MD simulation protocol

The CG dimer simulations of HP1α and pHP1α were performed in LAMMPS^81^ for 3 μs using the recently developed HPS-Urry and HPS-PTM models.^42,43^ CG coexistence simulations were conducted using the HOOMD-Blue 2.9.7 software package,^82^ using the protocol proposed in our previous work.^83,84^ To simulate HP1α and pHP1α homodimers, the folded domains were constrained using the hoomd.md.constrain.rigid function.^44^ CD domains were treated as separate rigid bodies while the CSD-CSD domains were held as a single rigid body. The initial slab configuration (17nm x 17nm x 119nm) was prepared from 50 dimer chains using the HPS-Urry and the HPS-PTM models. In the coexistence slab simulations, 5μs NVT runs were conducted at 320K using a Langevin thermostat with a friction factor, γ = m_AA_/*τ*. Here m_AA_ is the mass of each amino acid bead, *τ* is the damping factor, which was set to 1000 ps. The time step is set to 10 fs. When calculating the density profile and contact map, 1 μs trajectory was skipped for the equilibration. In the peptide titration simulation, the number of dimers of HP1α were kept constant and the number of peptides were varied to account for different ratios of peptide to pHP1α: 0.25, 0.5, 1, 2, 3, 4, 5. Whereas, in the slab simulations of HP1α and dsDNA, we kept the number of HP1α dimers constant (set to 50), and varied the number of dsDNA chains such that the mole fraction of dsDNA within the system was set to 0.02 and 0.038.

### Materials

All buffering salts, agarose, LB agar Miller, LB broth Miller, ampicillin sodium salt, isopropyl-β-D-thiogalactopyranoside (IPTG), glycerol, HisPur™ Ni-NTA resins, Pierce™ protease inhibitor tablets, Coomassie brilliant blue R-250, guanidine hydrochloride, magnesium sulfate heptahydrate, zinc chloride, copper(II) sulfate pentahydrate, and sodium dodecyl sulfate were purchased from ThermoFisher Scientific. 5x Phusion HF buffer and Phusion™ High-Fidelity DNA polymerase were purchased from ThermoFisher Scientific. Additional Phusion DNA polymerase for preparing 205 base pair DNA at large scales was provided by Professor Kevin Corbett’s lab at UCSD. Deoxynucleotides (dNTPs), 1 kb DNA ladder, 100 bp DNA ladder, Monarch® plasmid miniprep kit, DH5α competent cells, Rosetta (DE3) competent cells, Casein Kinase II (CKII), and 10x protein kinase buffer were purchased from New England BioLabs. Biomiga MV Gel/PCR extraction kits were purchased from Biomiga. Tris-(carboxyethyl) phosphine hydrochloride (TCEP) and adenosine-5’-triphosphate (ATP) disodium trihydrate were purchased from GoldBio. Imidazole was purchased from Acros Organics. DNase I, 100x Kao vitamin and 100x MEM vitamin solutions were purchased from Sigma Aldrich. Magnesium chloride hexahydrate was purchased from EMD. Sodium molybdate(VI) dihydrate and cobalt(II) chloride hexahydrate were purchased from Acros Organics. Iron(II) sulfate heptahydrate and sodium azide were purchased from Alfa Aesar. Ammonium chloride (^15^N, 99%), D-glucose (U-^13^C6, 99%), and deuterium oxide (D, 99.9%) were purchased from Cambridge Isotope Laboratory Inc. 10x tris/glycine/SDS buffer and Precision Plus Protein All Blue standards was purchased from Bio-Rad. Bromophenol blue sodium salt was purchased from MP Biomedical Inc. NMR tubes (3 mm and 5 mm OD, 800 MHz grade) were purchased from Wilmad.

### Instruments

Reverse phase (RP) high-performance liquid chromatography (HPLC) purification was performed on a Waters HPLC system using XBridge Peptide BEH C18 semi-prep column (5 μm, 10 mm x 250 mm) or preparative column (10 μm, 19 mm x 250 mm). The semi-preparative and preparative HPLC purifications were performed at 5 and 20 mL/min flowrates, respectively. The HPLC purifications used milli-Q (MQ) water with 0.1% TFA (solvent A) and acetonitrile with 0.1% TFA (solvent B). All gradient ranges were performed for 37 minutes. Nuclear magnetic resonance (NMR) data were collected using an 800 MHz Avance Neo Bruker NMR spectrometer equipped with a TXO cryo-probe optimized for ^15^N detection. An Olympus CKX53 microscope was used to collect images of the liquid-liquid phase separation droplets. A Thermo Scientific ™ NanoDrop ™ One^C^ Microvolume UV-Vis Spectrophotometer was used to measure the UV-vis absorbance of the HP1α constructs and peptides.

### Properties of HP1α and peptide ligands

The molecular weight, extinction coefficients at 280 nm, and charge were calculated on *http://protcalc.sourceforge.net/* using the amino acid sequence of each HP1α construct and peptide. The pH option was set to 7.2 for the charge calculation. The extinction coefficient at 205 nm was calculated on *https://spin.niddk.nih.gov/clore/*. The properties of the HP1α constructs and peptides are shown in **Table S3**.

### HP1α constructs

Wild-type human heterochromatin protein 1α (HP1α) was cloned into a pET vector to yield a His_6_-TEV-HP1α construct as described previously.^67^ This plasmid was used to clone all HP1α constructs described below. Cloning was performed using the NEBuilder® HiFi strategy following the protocol provided by the manufacturer. Plasmids were transformed into DH5α competent cells, the cells were plated on LB agar supplemented with ampicillin (100 μg/mL) and left overnight at 37 °C. A colony was selected and grown in 5 mL of LB broth with ampicillin (100 μg/mL) overnight at 37 °C with 220 rpm agitation. Plasmids were purified using the Monarch® plasmid miniprep kit (New England Biolabs) and sequenced to confirm the desired cloning result (Azenta/Genewiz). All HP1α constructs have an N-terminus starting sequence of MKSSHHHHHHENLYFQ, which was cleaved by TEV protease during purification. Due to the N-terminal TEV cleavage site, all HP1α constructs start with a serine residue at the N-terminus. The amino acid sequences of the HP1α constructs are shown below in bold.

HP1α, residues 1-191 (wild-type):

**SGKKTKRTADSSSSEDEEEYVVEKVLDRRVVKGQVEYLLKWKGFSEEHNTWEPEKNLDCPELISEFMKKYKKMKEGENNKPREKSESNKRKSNFSNSADDIKSKKKREQSNDIARGFERGLEPEKIIGATDSCGDLMFLMKWKDTDEADLVLAKEANVKCPQIVIAFYEERLTWHAYPEDAENKEKETAKS**

HP1α-W174A, residues 1-191:

**SGKKTKRTADSSSSEDEEEYVVEKVLDRRVVKGQVEYLLKWKGFSEEHNTWEPEKNLDCPELISEFMKKYKKMKEGENNKPREKSESNKRKSNFSNSADDIKSKKKREQSNDIARGFERGLEPEKIIGATDSCGDLMFLMKWKDTDEADLVLAKEANVKCPQIVIAFYEERLTAHAYPEDAENKEKETAKS**

HP1α chromoshadow domain (HP1α-CSD), residues 112-176:

S**DIARGFERGLEPEKIIGATDSCGDLMFLMKWKDTDEADLVLAKEANVKCPQIVIAFYEERLTWHA**

HP1α chromoshadow domain with C-terminus extension (HP1α-CSDCTE), residues 110-191:

S**SNDIARGFERGLEPEKIIGATDSCGDLMFLMKWKDTDEADLVLAKEANVKCPQIVIAFYEERLTWHAYPEDAENKEKETAKS**

HP1α-CSDCTE-W174A, residues 110-191:

**SSNDIARGFERGLEPEKIIGATDSCGDLMFLMKWKDTDEADLVLAKEANVKCPQIVIAFYEERLTAHAYPEDAENKEKETAKS**

### 205 bp DNA construct

The 205 bp double stranded DNA construct used for LLPS has the following sequence: 5’-

**CTGGAGAATCCCGGTGCCGAGGCTGCAGAATTGGTCGTAGACAGCTCTAGCACCGCTTAAACGCACGTACGCGCTGTCCCCCGCGTTTTAACCGCCAAGGGGATTACTCCCTAGTCTCCAGGCACGTGTCAGATATATACATCCTGTGCATGTATTGAACAGCGACCTTGCCGGTGCCAGTCGGATAGTGTTCCGAGCTCCCTGT**

-3’.

The 205 bp DNA was amplified using polymerase chain reaction (PCR) with the forward primer 5’-CTGGAGAATCCCGGTGCCGAGGCTGCAGAATTG-3’ and the reverse primer 5’-ACAGGGAGCTCGGAACACTATCCGACTGGCACCGGC-3’, each added at 0.5 μM concentration. 5x Phusion HF buffer was added equal to one-fifth of the total PCR volume along with Phusion DNA polymerase (generously provided by Dr. Kevin Corbett) and dNTPs at 600 μM final concentration. PCR was performed with the following thermal cycle: (1) initial denaturing temperature was 98 °C for 30 seconds, (2) denaturing temperature was 98 °C for 10 seconds, (3) annealing temperature was 60.0 °C, (4) extension time was 20 seconds at 72 °C, (5) steps 2-4 were repeated for 35 cycles, (6) final extension time was 5 minutes at 72 °C. The PCR product was purified using a Biomiga PCR cleaning kit following the procedure provided by the manufacturer. The 205 bp DNA was stored in 20 mM HEPES, pH 7.2, and 1 mM TCEP. The purity of the final DNA product was confirmed on a 5% TBE acrylamide gel stained with ethidium bromide.

### Expression, purification, and refolding of HP1α constructs

All HP1α constructs were expressed in Rosetta (DE3) competent cells cultured in Luria-Bertani (LB) broth media. ^15^N- and/or ^13^C-isotopically labeled HP1α constructs for NMR spectroscopy were prepared using minimal media (M9) supplemented with ammonium chloride (^15^N, 99 %) and glucose (U-^13^C6, 99 %). The expression, purification and refolding protocols were adapted from the following references ^67 31^.

#### Expression of HP1α Constructs

The DNA plasmid containing the desired HP1α construct was transformed into Rosetta cells following standard transformation protocols. An isolated colony was selected for the starter culture and grown in 20 mL of LB broth with ampicillin (100 μg/mL final concentration) at 37 °C with 220 rpm agitation. 10 mL of the starter culture were spun down, and the cell pellet was resuspended in 10 mL of fresh LB broth. The resuspension was added to 1 L of LB broth with ampicillin (100 μg/mL final concentration) and cells were cultured at 37 °C with 220 rpm agitation until the OD600 reached 0.6-0.8. At this point, IPTG was added into the culture to a final concentration of 0.5 mM to induce the expression of HP1α. The induced culture was grown at 18 °C with 220 rpm agitation overnight.

#### Purification of HP1α Constructs

After overnight growth, the culture was spun down at 5,000 xg for 25 minutes. The supernatant was discarded, and the cell pellet was resuspended in 25 mL of lysis buffer (20 mM HEPES, pH 7.2, 300 mM KCl, 10% glycerol, and one Pierce protease inhibitor tablet). The sample containing HP1α was kept on ice throughout the purification process. While on ice, the cells were lysed via sonication for 10 minutes with a 30 second on/off cycle at 4 °C.

After sonication, the lysate was centrifuged at 30,000 xg for 20 minutes. Approximately 4 mL of Ni-NTA resin slurry was added into the supernatant along with 8 mg of DNase I and MgCl_2_ at 10 mM final concentration. The mixture, consisting of the supernatant and Ni-NTA resin, was rotated at 4 °C for at least one hour. The mixture was then transferred into a column, the Ni-NTA resin was left to settle for ∼ 10 min, and the supernatant was collected by elution. The Ni-NTA resin was washed two times using 25 mL of wash buffer (20 mM HEPES, pH 7.2, 300 mM KCl, 10% glycerol, 40 mM imidazole). To elute HP1α, 35 mL of buffer (20 mM HEPES, pH 7.2, 300 mM KCl, 400 mM imidazole) were added to the resin and 35 mL elution volume was collected. After elution, TCEP (0.5 mM final concentration) was added to the fractions containing HP1α to prevent disulfide bond formation. The 6xHis-tag on HP1α was removed by adding 800 μg of TEV protease to the combined HP1α elution fractions. Concurrent with TEV cleavage, the HP1α solution was dialyzed against 20 mM HEPES, pH 7.2, 300 mM KCl, 0.5 mM TCEP at 4 °C overnight. Sample purity and 6xHis-tag cleavage were confirmed on a 15% acrylamide SDS PAGE gel stained with Coomassie. The HP1α sample was then prepared for HPLC purification by adding solid guanidine hydrochloride (GdHCl) to a final concentration of 6M, and pH was adjusted to ∼ 2 – 4 using trifluoroacetic acid (TFA). The solution was filtered through a 0.45 μm syringe filter (HPLC grade) and loaded onto a preparative, reverse phase C18 HPLC column for HPLC purification. HP1α was eluted using a gradient of 30-45 % solvent B over 37 min. The collected fractions were analyzed on a 15% acrylamide SDS PAGE gel. Pure fractions were pooled, lyophilized, and stored at – 80 °C. Final purity was assessed by LC-ESI-TOF-MS (**Fig. S2**).

#### Refolding of HP1α Constructs

Lyophilized HP1α was dissolved into 25 mL of buffer (20 mM HEPES, pH 7.2, 300 mM KCl, 6M GdHCl, 1 mM TCEP). The solution was transferred into a 10,000 MWCO SnakeSkin® dialysis tubing and dialyzed against 1 L of buffer (20 mM HEPES, pH 7.2, 1 M GdHCl, 300 mM KCl, 1 mM TCEP) at 4 °C for ∼ 6-8 hours with stirring. The solution was then dialyzed against 1 L of buffer (20 mM HEPES, pH 7.2, 300 mM KCl, 1 mM TCEP) at 4 °C overnight with stirring, followed by dialysis against 1 L of fresh buffer for additional 6-8 hours. After refolding, the solution was concentrated to a volume of 200-500 μL, and 1 mL of fresh buffer (20 mM HEPES, pH 7.2, 300 mM KCl, 1 mM TCEP) was added. The concentration/buffer exchange steps were repeated three times to remove any residual GdHCl left after dialysis. The typical yield for the full-length HP1α constructs was 15-20 mg per liter of LB broth expression. For the HP1α-CSD/CSDCTE construct, the typical yield was 5-10 mg per liter of LB broth expression.

### Peptide purification

The peptides were derived from the following HP1α protein-binding partners: lamin B receptor (LBR), chromatin assembly factor 1 (CAF-1), shugoshin (Sgo1), histone H3 (H3), and histone H4 (H4) proteins (**Table S3**). Sgo1 was purchased from ABclonal Technology (Woburn, MA) at 95% purity. LBR, CAF-1 and H3 were purchased from ABclonal Technology (Woburn, MA) at crude level of purity. H4 was purchased from InnoPep (San Diego, CA) at crude level of purity. Crude peptide samples were purified using HPLC, and the final purity was confirmed using LC-ESI-TOF-MS. Purifications were carried out on a semi-preparative, reverse phase HPLC column with a 37 min gradient with the following solvent ranges: LBR (20-50% solvent B), CAF-1 (30-70% solvent B), H3 (10-60% solvent B), and H4 (0-50% solvent B). Pure peptide fractions were lyophilized and stored at -20 °C. For LLPS assays, the peptides were dissolved in buffer containing 20 mM HEPES, pH 7.2, and 1 mM TCEP when needed. The concentration of Sgo1 and H3 was determined using absorbance at 280 nm. The concentration of LBR, CAF-1, and H4 was estimated using absorbance at 205 nm.^85^ (See **Table S3**)

### Analytical ultracentrifugation (AUC) of HP1α constructs

AUC-Sedimentation Velocity (AUC-SV) experiments were performed with the full-length HP1α and HP1α-CSD, separately, in a ProteomeLab XL-I (BeckmanCoulter) analytical ultracentrifuge using absorbance at 280 nm for detection. Protein samples were loaded in 2-channel cells with sapphire windows and centrifuged in An-50 Ti 8-place rotor at 40,000 rpm, 20 °C for 20 hours. Data were analyzed using the Sedfit software (P. Schuck, NIH/NIBIB). Detection cut-off was 1% of the total protein amount in the samples.

### In-vitro phosphorylation of HP1α constructs

Refolded HP1α was phosphorylated *in vitro* with Casein Kinase II (CK2) purchased from New England Biolabs, Inc. Typically, 0.3 mL of 25 mg/mL HP1α solution was diluted in 1X protein kinase buffer (supplied by the manufacturer). The solution also contained ATP at a mole ratio of 28 ATP molecules to one HP1α monomer. To initiate phosphorylation, 5 μL of CK2 were added to the sample and the solution was incubated at 30 °C overnight. After incubation, KCl was added to a final concentration of 300 mM. The mixture was concentrated to ∼ 100 μL and buffer exchanged using 20 mM HEPES, pH 7.2, 300 mM KCl, and 1 mM TCEP, using a 10,000 MWCO concentrator to remove ATP from the solution. The buffer exchange was performed at least 5 times or until the A260/280 ratio was below 0.6. Phosphorylation was confirmed by 15% SDS PAGE analysis, LC-MS ESI TOF, and 5% TBE native gel electrophoresis analysis (**Fig. S2 and Fig. S12a**). LC-MS/MS analysis was performed to confirm the identity of the phosphorylated sites. For this purpose, phosphorylated HP1α was analyzed on a 15% acrylamide SDS PAGE gel, the relevant band was excised, the protein was digested with trypsin and subjected to a 45 min LC run followed by MS/MS analysis (**Table S1**). Data were analyzed with MaxQuant using the standard setup and the following parameters: fixed parameters (Carbamidomethyl (C)), variable parameters (Oxidation (M)), acetyl (protein N-term), phospho (STY)), and digestion mode (Trypsin/P), along with a fasta file of the HP1α sequence.^86^

### Peptide binding assays

To verify that HP1α was properly folded and functional, we performed a binding assay with Sgo1 where binding was verified by gel electrophoresis (**Fig. S12a**). HP1α and pHP1α (25 μM) were incubated without and with 25 μM Sgo1 in buffer (20 mM HEPES, pH 7.2, 300 mM KCl, 1 mM TCEP, 10% glycerol) for at least 20 minutes at room temperature. For each sample, 10 μL was loaded onto a 5% TBE native gel and analyzed by native gel electrophoresis.

### NMR titration experiments

#### Sample preparation

All NMR experiments were performed in 20 mM HEPES, pH 7.2, 75 mM KCl, 1 mM TCEP, 0.05% NaN_3_. Samples contained 400 μM of HP1α-CSD in a 5 mm NMR tube (500 μL volume) for chemical shift assignments, or 100 μM of HP1α-CSD in a 3 mm NMR tube (150 μL) for titration experiments. Peptide concentrations used for the titration experiments were 13, 25, 50, 100, and 200 μM, respectively. For experiments with the HP1α-CSDCTE, 70 μM protein was prepared into a 3 mm NMR tube (150 μL volume). Peptide concentrations in this case were 9, 18, 35, 70, and 140 μM, respectively. For the W174A mutant, 50 μM HP1α-CSDCTE-W174A was used along with 50 μM of peptide resulting in a 2 to 1 ratio of peptide to homodimer.

#### NMR experiments

The NMR data were collected using an 800 MHz Avance Neo Bruker NMR spectrometer equipped with a TXO cryo-probe optimized for ^15^N detection. Chemical shifts were referenced relative to the spectrometer frequency, with the water resonance at 4.7 ppm. Pulses were calibrated using the standard Topspin protocol. Chemical shift assignments were performed using standard 2D ^1^H-^15^N HSQC, 3D HNCACB, and 3D CBCA(CO)NH experiments. HSQC experiments were performed with the pulse program fhsqcf3gpph,^87^ with 16 scans and a relaxation delay of 1 sec. For the NMR titration data sets, 48 scans were used. The following parameters were set for the ^1^H dimension: 13.58 ppm spectral width, frequency offset of 4.723 ppm, 2048 points and DQD acquisition mode. The following parameters were used for the ^15^N dimension: 36.00 ppm spectral width, frequency offset of 120 ppm, 128 points and States-TPPI acquisition mode. The 3D HNCACB experiment was performed with the pulse sequence hncacbgpwg3d^88,89^ with 32 scans and a relaxation delay of 1 sec. The 3D CBCA(CO)NH experiment was performed with the pulse sequence cbcaconhgpwg3d,^89,90^ 28 scans and a relaxation delay of 1 sec. For the ^1^H dimension in both experiments, the following parameters were used: 14.20 ppm spectral width, frequency offset of 4.734 ppm, 2048 points and DQD acquisition mode. The following parameters were used for the ^15^N dimension: 30.00 ppm spectral width, frequency offset of 118 ppm, 64 points and States-TPPI acquisition mode. The following parameters were used for the ^13^C dimension: 64.00 ppm spectral width, frequency offset of 40.50 ppm, 108 points and States-TPPI acquisition mode. Data were processed using NMRPipe^91^ and analyzed in NMRFAM-SPARKY^92^ (Sparky). Backbone chemical shifts (amide proton (^1^H), amide nitrogen (^15^N), alpha carbon (Cα), and beta carbon (Cβ)) were assigned for all residues except the N-terminal serine (present due to TEV cleavage), Pro 122, Pro 161, and Asp 112 (Cα and Cβ chemical shift assigned).

### Liquid-liquid phase separation (LLPS) of HP1α

#### LLPS of pHP1α with peptides

We performed two types of experiments. In the first type of experiments, we varied the concentration of pHP1α or pHP1α-W174A (25, 50, 100, 150, 200, and 250 μM) and held the peptide concentration constant (75 μM). Note that the pHP1α or pHP1α-W174A numbers reflect the concentration of the monomeric protein. In the second type of experiments, we kept the concentration of protein constant, and varied the concentration of the peptide. In this case, the concentration of pHP1α was 200 μM, and ratios of peptide to homodimer of 0, 0.08, 0.12, 0.24, 0.48, 0.71, 0.95, 1.42, 1.90, and 2.85 were used. For experiments with pHP1α-W174A, the concentration of protein was 200 μM, and peptide concentrations of 25 and 100 μM of peptide were used. LLPS experiments were performed in 20 mM HEPES, pH 7.2, 75 mM KCl, and 1 mM TCEP. After all the necessary components were added, the samples were mixed by pipetting up and down a few times, then incubated on ice for at least 20 minutes before data collection. Samples were kept on ice or at 4°C throughout the course of the experiments. A no LLPS control was prepared by adding pHP1α to a buffer containing 20 mM HEPES, pH 7.2, 300 mM KCl, and 1 mM TCEP.

#### LLPS of HP1α with DNA and peptides

We performed two types of experiments. In the first case, we varied the concentration of HP1α (25, 50, 100, 150, 200, and 250 μM), and kept the concentration of 205 bp DNA (1.5 μM) and peptide constant (75 μM). In the second case, we kept the concentration of HP1α (100 μM) and peptide constant (75 μM), and varied the concentration of DNA (0.5, 1.0, 1.5, 2.0, 2.5, 3.0, 3.5, and 4.0 μM). For LLPS with HP1α-W174A, we used 100 μM of protein, 75 μM of peptide, and varied the DNA concentration (0.5, 1.5, and 2.5 μM). LLPS experiments were performed in 20 mM HEPES, pH 7.2, 75 mM KCl, and 1 mM TCEP. After all the necessary components were added, the samples were mixed by pipetting up and down a few times, then incubated on ice for at least 20 minutes before data collection. Samples were kept on ice or at 4°C throughout the course of the experiments. A no LLPS control was prepared by adding HP1α to a buffer containing 20 mM HEPES, pH 7.2, 75 mM KCl, and 1 mM TCEP, without the presence of DNA.

#### Microscopy experiments

An Olympus CKX53 microscope was used to image the HP1α liquid droplets. Before loading the sample onto the microscopy glass slide, the solution was mixed by pipetting up and down a few times, and 6-8 μL were loaded onto the slide. Images were acquired within 1-2 minutes. For fluorescence imaging of pHP1α, 0.6 μM of pHP1α conjugated with a Cy3 dye (pHP1α-Cy3) was introduced into the samples. For imaging of DNA, we used 0.1 μM of YOYO-1 dye.

#### Absorbance measurements

The absorbance at 280 nm (A280) and the ratio between the absorbance at 260 nm and 280 nm (A260/A280) of the supernatant phase were measured using a Thermo Scientific ™ NanoDrop ™ One^C^ Microvolume UV-Vis Spectrophotometer. For LLPS samples, the condense (liquid droplet) phase and dilute (supernatant) phase were separated by centrifugation at 200 xg for 10 minutes at 4°C. 2 μL of the supernatant was used for each measurement. For pHP1α samples, the A280 measurements were converted directly into the molar concentration of the monomeric protein using the extinction coefficients of the wild type and the W174A mutant HP1α constructs (**Table S3**). For samples that contained DNA, we used the A280 automated correction built into the Acclaro™ software of the NanoDrop that takes into account the A260/A280 ratio of the sample. The corrected A280 was then converted into molar concentration.

## Acknowledgments

We thank Dr. Xuemei Huang for help with NMR experiments, Dr. Andrey Bobkov for collecting the AUC data, and Dr. Majid Ghassemian and Dr. Yongxuan Su for help with mass spec data collection and analysis. This work was supported by NIH grants R35GM138382 (G.T.D., C.H.) and R01GM136917 (J.M., T.M.P, N.J., U.K., A.R.). Y.C.K. is supported by the Office of Naval Research via the U.S. Naval Research Laboratory base program. B.E.A. is supported by the NIH Molecular Biophysics Training Grant T32 GM008326. AUC was performed at the protein analysis facility at the Sanford Burnham Prebys Medical Discovery Institute supported by NIH P30 CA030199.

## Author contributions

CH, TMP, JM, and GTD conceived and designed research. CH and BEA designed and conducted wet lab experiments. TMP and NJ performed computer simulations of HP1α, and UK conducted the simulations with DNA. AR and YCK provided the computational tools needed to setup the coarse-grained simulations. JM and GTD supervised research. CH, TMP, JM, and GTD wrote the manuscript with help from other authors.

## Competing interests

The authors declare that they have no competing interests.

## Notes

### Competing Interest Statement

The authors have declared no competing interest.

