## Supporting information for "Molecular interactions underlying the phase separation of HP1α: Role of phosphorylation, ligand and nucleic acid binding"

#### Supporting information figures

**(a)**

HP1α Sedimentation Velocity AUC

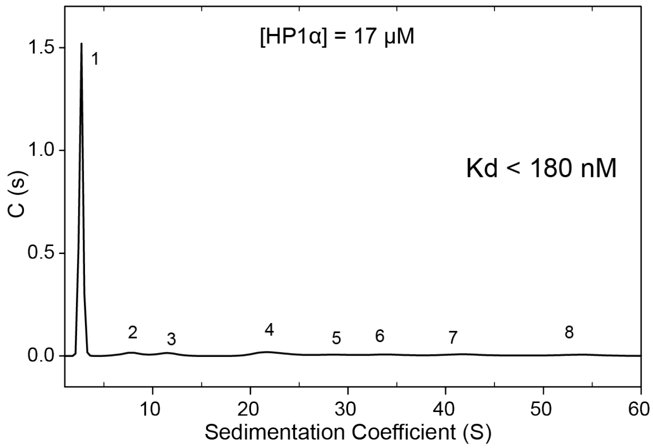


| Apparent Molecular Weight of Each Peak | | | | |
|---|---|---|---|---|
| **Peak #** | **S** | **S (20,w)** | **% of total protein** | **Ap. Mw (kDa)** |
| 1 | 2.688 | 2.837 | 64.9 | 39 |
| 2 | 7.805 | 8.236 | 3.8 | 193 |
| 3 | 11.556 | 12.195 | 3.3 | 348 |
| 4 | 22.382 | 23.618 | 7.3 | 939 |
| 5 | 28.753 | 30.342 | 2.5 | 1,367 |
| 6 | 34.242 | 36.134 | 3.2 | 1,777 |
| 7 | 42.265 | 44.600 | 4.4 | 2,437 |
| 8 | 53.256 | 56.198 | 3.7 | 3,447 |

**(b)**

HP1α-CSD Sedimentation Velocity AUC

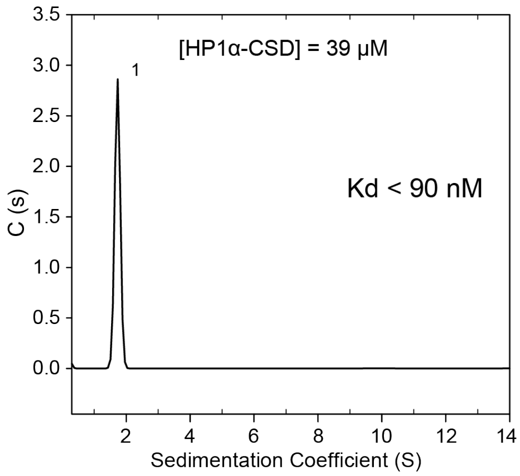


| Apparent Molecular Weight of Each Peak | | | | |
|---|---|---|---|---|
| **Peak #** | **S** | **S (20,w)** | **% of total protein** | **Ap. Mw (kDa)** |
| 1 | 1.729 | 1.778 | - | 15.4 |

**Figure S1.** Analytical ultracentrifugation (AUC) of HP1α and HP1α-CSD using the following buffer conditions: 20 mM HEPES, pH 7.2, 75 mM KCl, and 1 mM TCEP. Representative sedimentation velocity-AUC (SV-AUC) experiments of **(a)** HP1α at 17 μM and **(b)** HP1α-CSD at 39 μM. The tables show the apparent molecular weight and relative amount of protein calculated based on the sedimentation coefficient (S) of the detected species. The dissociation constant (Kd) of the HP1α homodimer and the HP1α-CSD homodimer were estimated to be less than 180 nM and 90 nM, respectively.

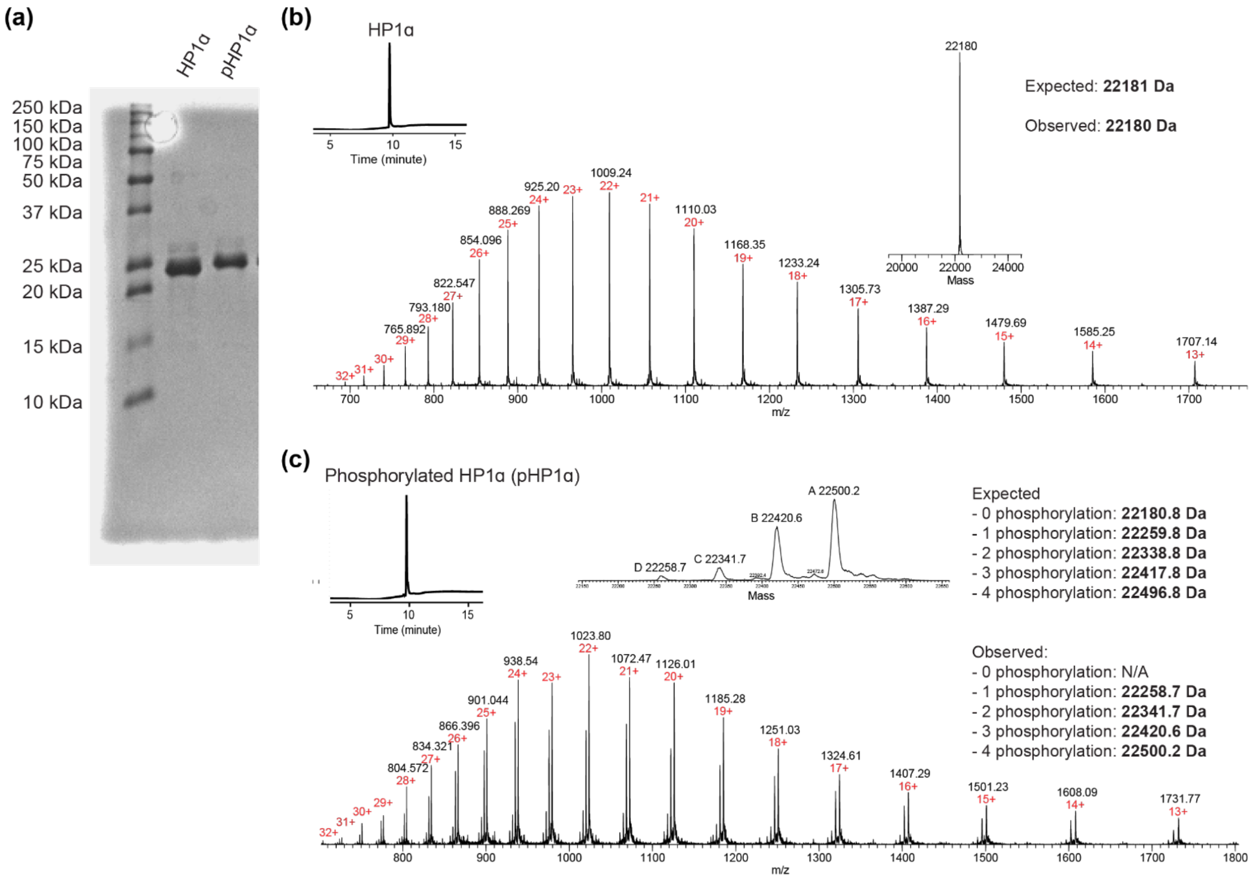


**Figure S2.** Analysis of purified HP1α and pHP1α. **(a)** 15% acrylamide SDS PAGE gel electrophoresis analysis. **(b)** LC-MS analysis of purified HP1α. The RP-HPLC trace is shown in the top left corner. **(c)** LC-MS analysis of purified pHP1α. The MS analysis indicates the presence of pHP1α containing on average one, two, three or four phosphorylation sites. The RP-HPLC trace is shown in the top left corner.

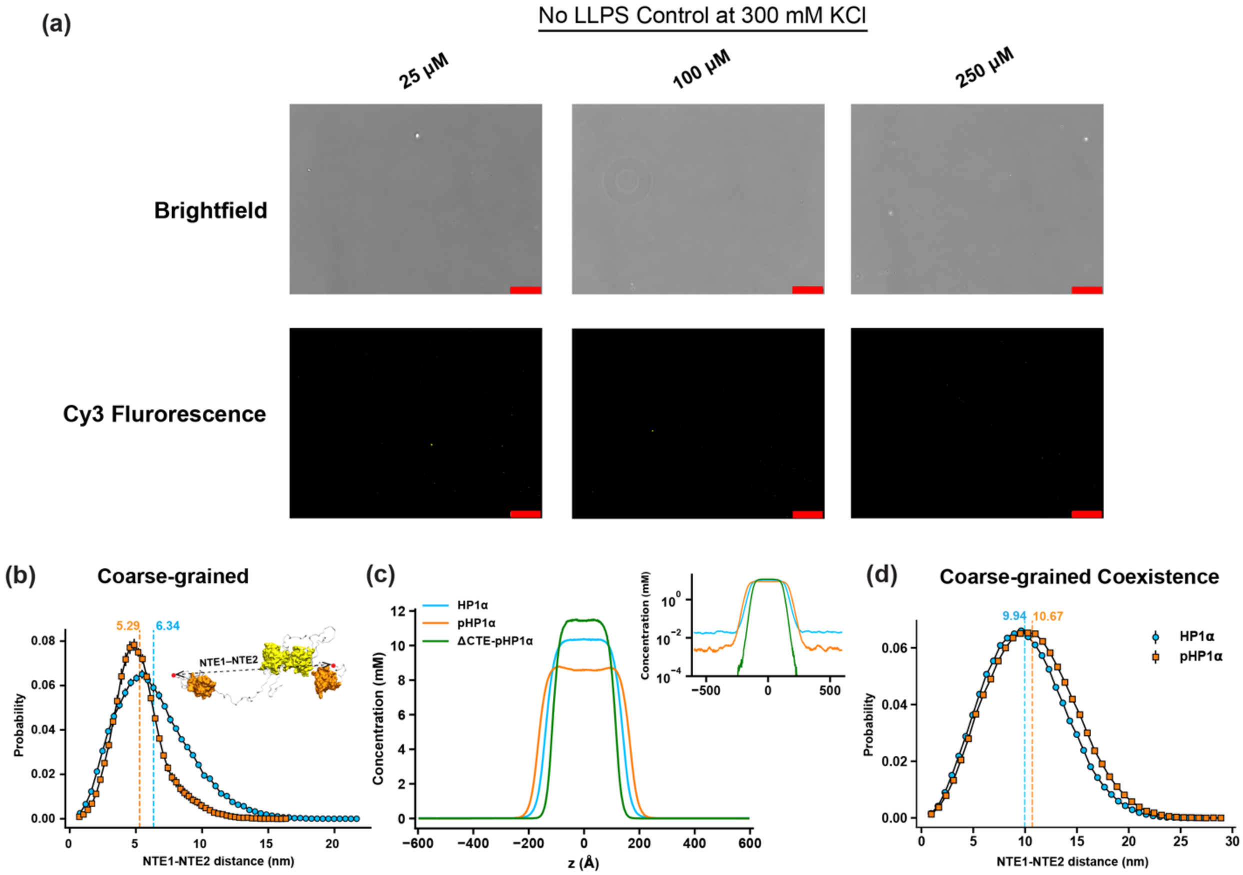


**Figure S3. (a)** Brightfield and Cy3 fluorescence images of the no LLPS control of pHP1α in the following buffer: 20 mM HEPES, pH 7.2, 300 mM KCl, and 1 mM TCEP. The red scale bar represents 100 μm. **(b)** Distribution distance between NTE1 and NTE2 of HP1α and pHP1α obtained from CG single homodimer simulations. Dashed lines represent mean values of each distribution. Errors are estimated using block averages with five blocks. **(c)** Density profiles of HP1α, pHP1α, and pHP1α with a deletion of the last 14 residues of the CTE (ΔCTE- pHP1α). **(d)** Distribution distance between NTE1 and NTE2 of HP1α and pHP1α obtained from CG co-existence simulations.

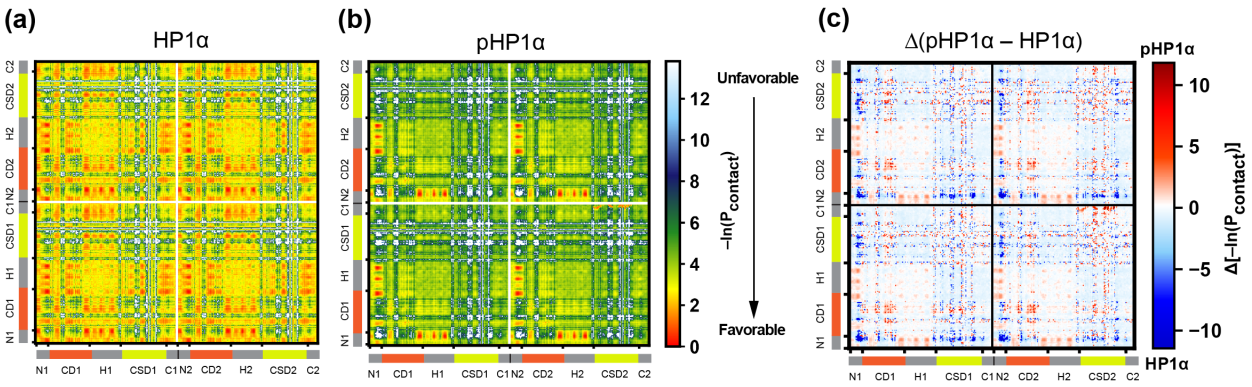


**Figure S4. (a, b)** Intermolecular contact maps of HP1α and pHP1α in CG coexistence simulations. **(c)** Contact map difference between HP1α and pHP1α. Blue and red color schemes show the preferential interactions towards HP1α and pHP1α, respectively.

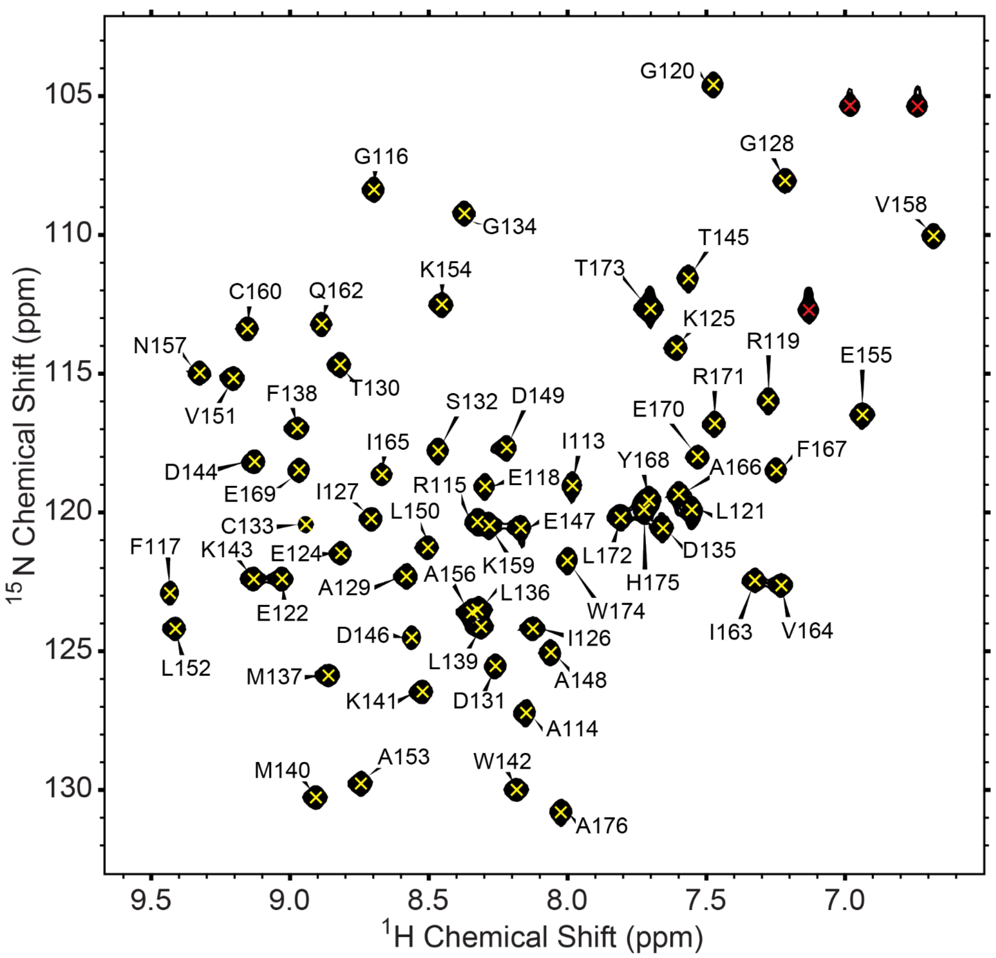


**Figure S5.** 2D $^{1}$H-$^{15}$N HSQC NMR spectrum of HP1α-CSD. The cross peaks are labeled with their corresponding residue. The cross peaks marked with a red 'x' are from side chains.

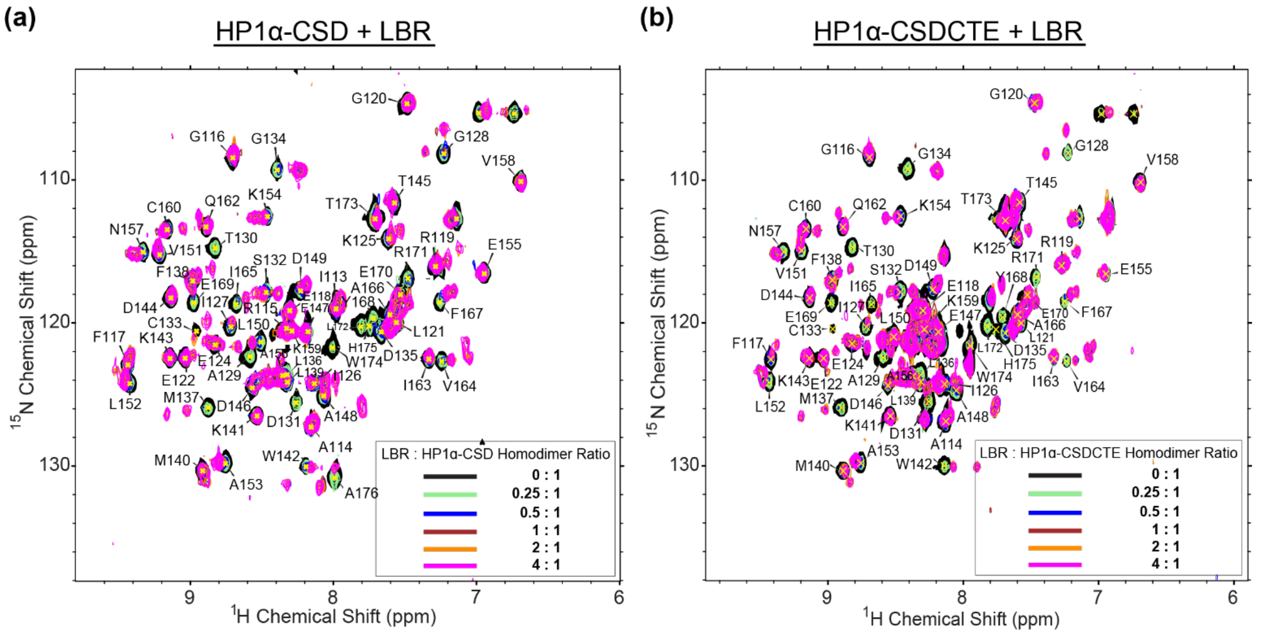


**Figure S6.** Overlay of 2D $^{1}$H-$^{15}$N HSQC NMR spectra of **(a)** HP1α-CSD and **(b)** HP1α-CSDCTE with LBR peptide present at different concentrations. The cross peaks are assigned based on the HP1α-CSD control.

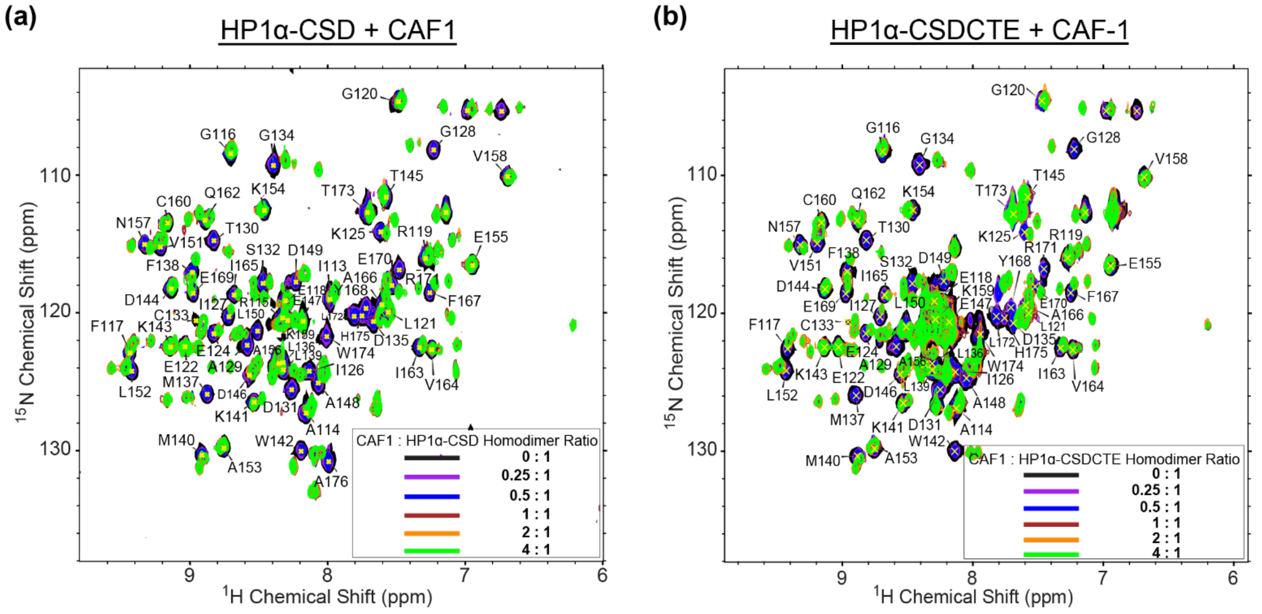


**Figure S7.** Overlay of 2D $^{1}$H-$^{15}$N HSQC NMR spectra of **(a)** HP1α-CSD and **(b)** HP1α-CSDCTE with CAF1 peptide present at different concentrations. The cross peaks are assigned based on the HP1α-CSD control.

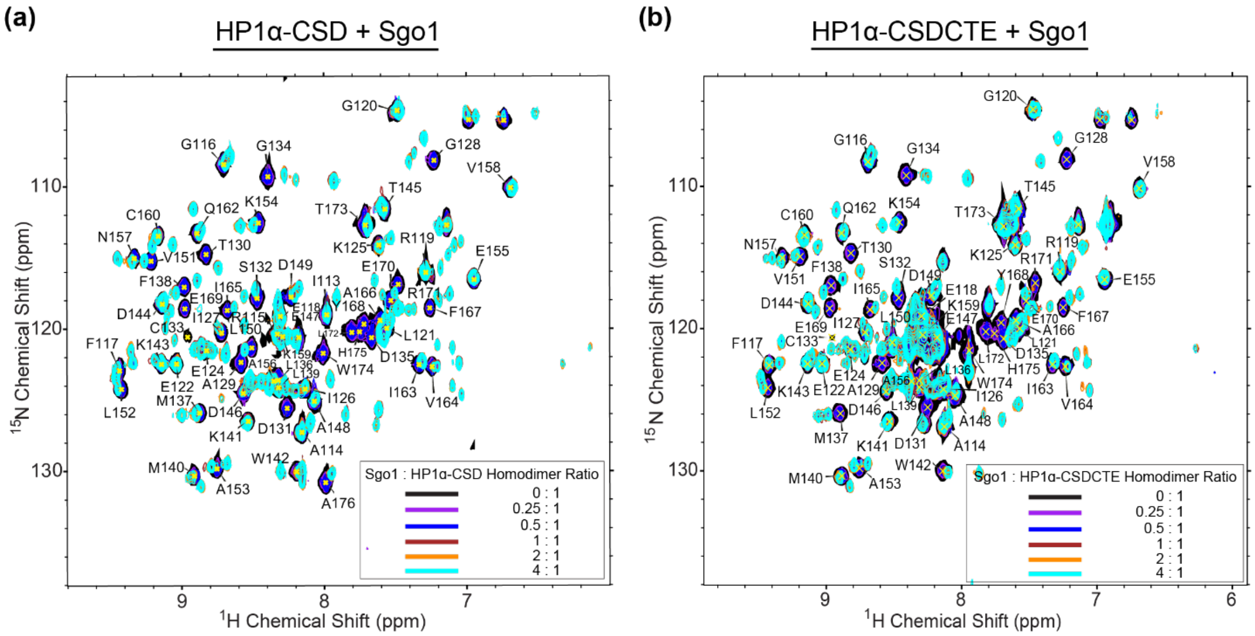


**Figure S8.** Overlay of 2D $^{1}$H-$^{15}$N HSQC NMR spectra of **(a)** HP1α-CSD and **(b)** HP1α-CSDCTE with Sgo1 peptide present at different concentrations. The cross peaks are assigned based on the HP1α-CSD control.

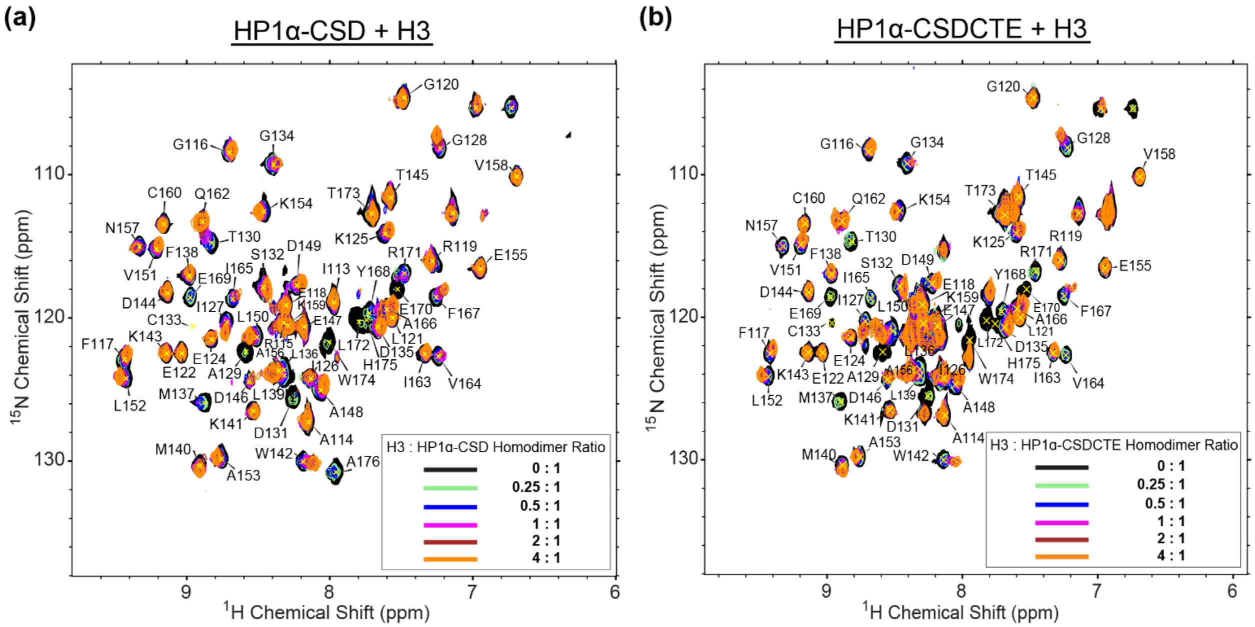


**Figure S9.** Overlay of 2D $^{1}$H-$^{15}$N HSQC NMR spectra of **(a)** HP1α-CSD and **(b)** HP1α-CSDCTE with H3 peptide present at different concentrations. The cross peaks are assigned based on the HP1α-CSD control.

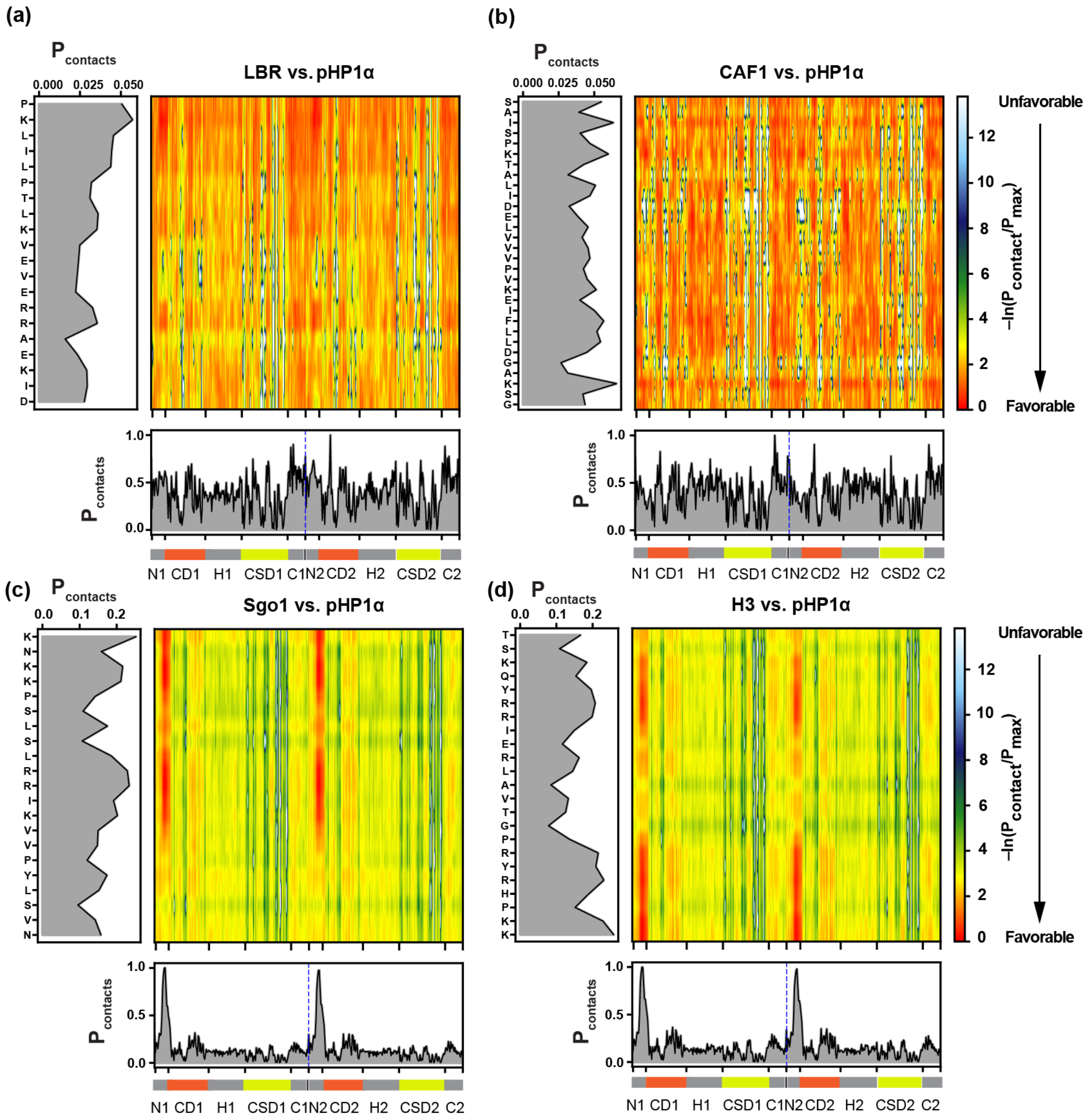


**Figure S10.** Intermolecular contacts between peptides and pHP1α in coexistence simulations. One-dimensional plots show the total contact propensity as a function of protein residue (HP1α at the bottom and peptide on the left) per frame, averaged over the entire trajectory.

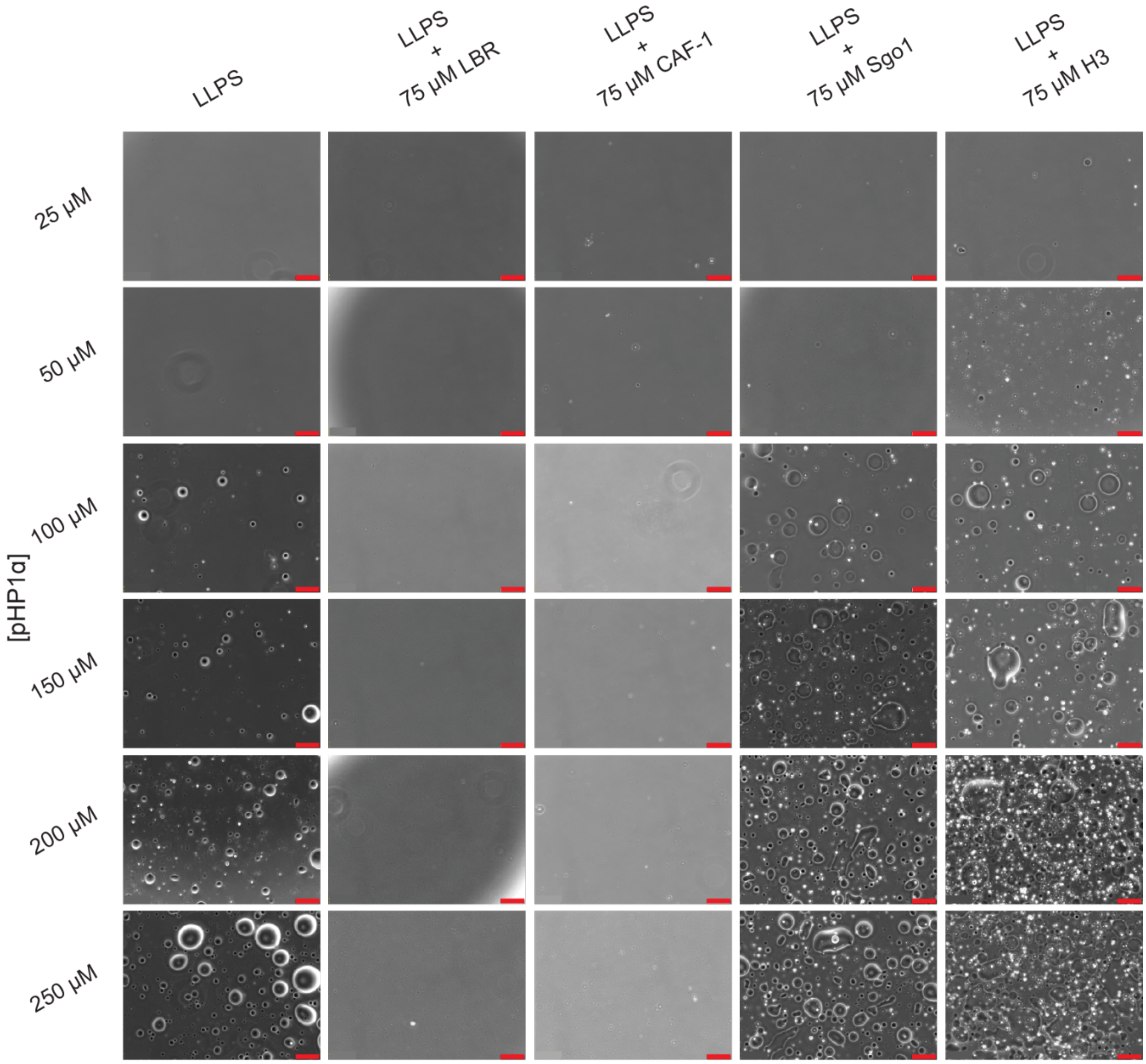


**Figure S11.** Brightfield microscopy images of pHP1α LLPS with and without 75 μM peptide as indicated on top of each column. The red scale bar represents 100 μm.

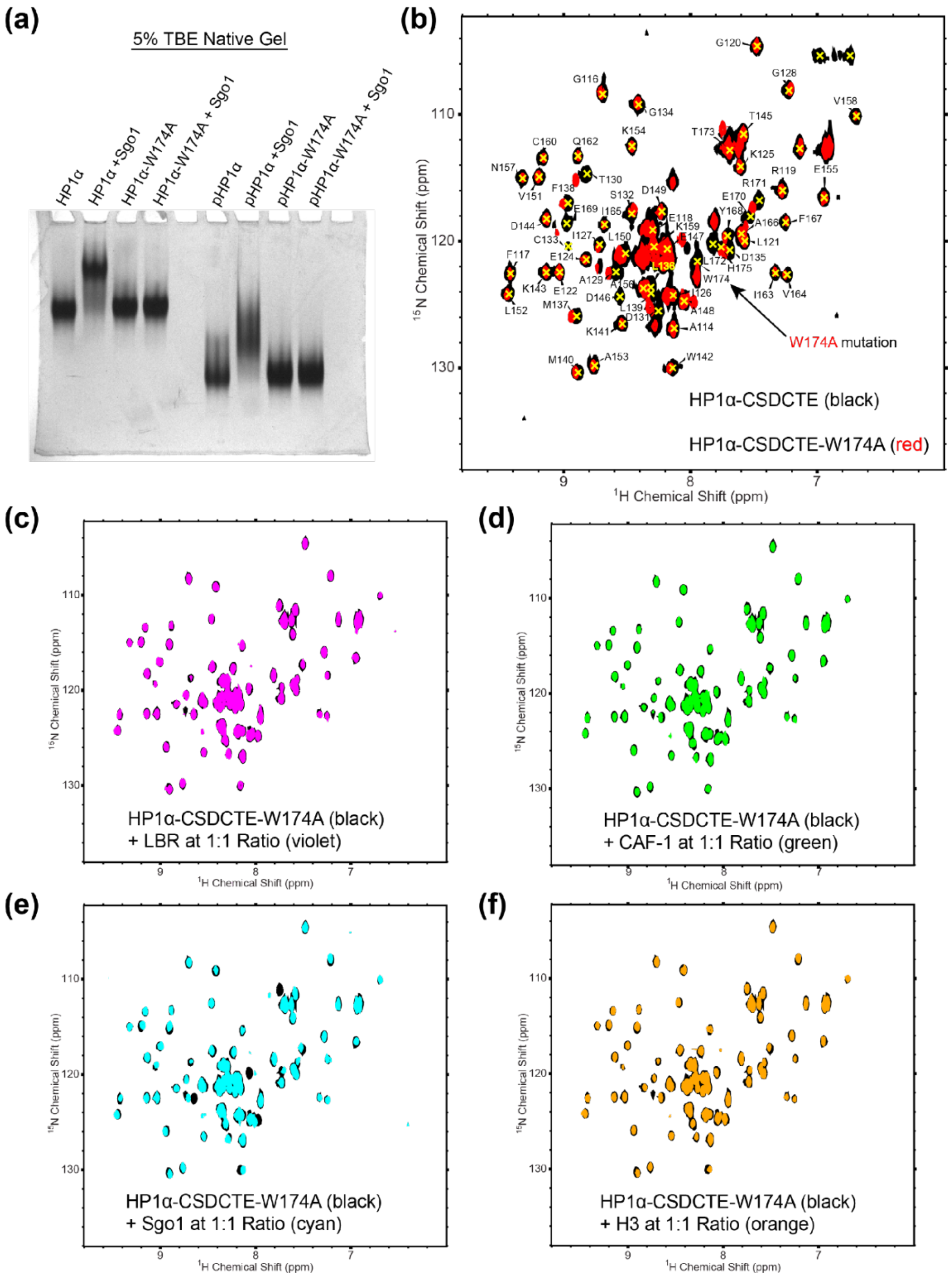


**Figure S12.** Role of Trp174 in ligand binding to the HP1α homodimer binding interface. **(a)** Interactions of the Sgo1 peptide with HP1α, HP1α-W174A, pHP1α, and pHP1α-W174A as analyzed by native PAGE. **(b)** Overlay of the 2D $^{1}$H-$^{15}$N HSQC NMR spectra of HP1α-CSDCTE (black) and HP1α-CSDCTE-W174A (red). The arrow indicates the location of the W174 peak in the NMR spectrum. The chemical shift assignments are based on HP1α-CSD. (**c-f**) Overlay of the 2D $^{1}$H-$^{15}$N HSQC NMR spectra of HP1α-CSDCTE-W174A without peptide (black) and with **(c)** LBR, **(d)** CAF-1, **(e)** Sgo1, or **(f)** H3.

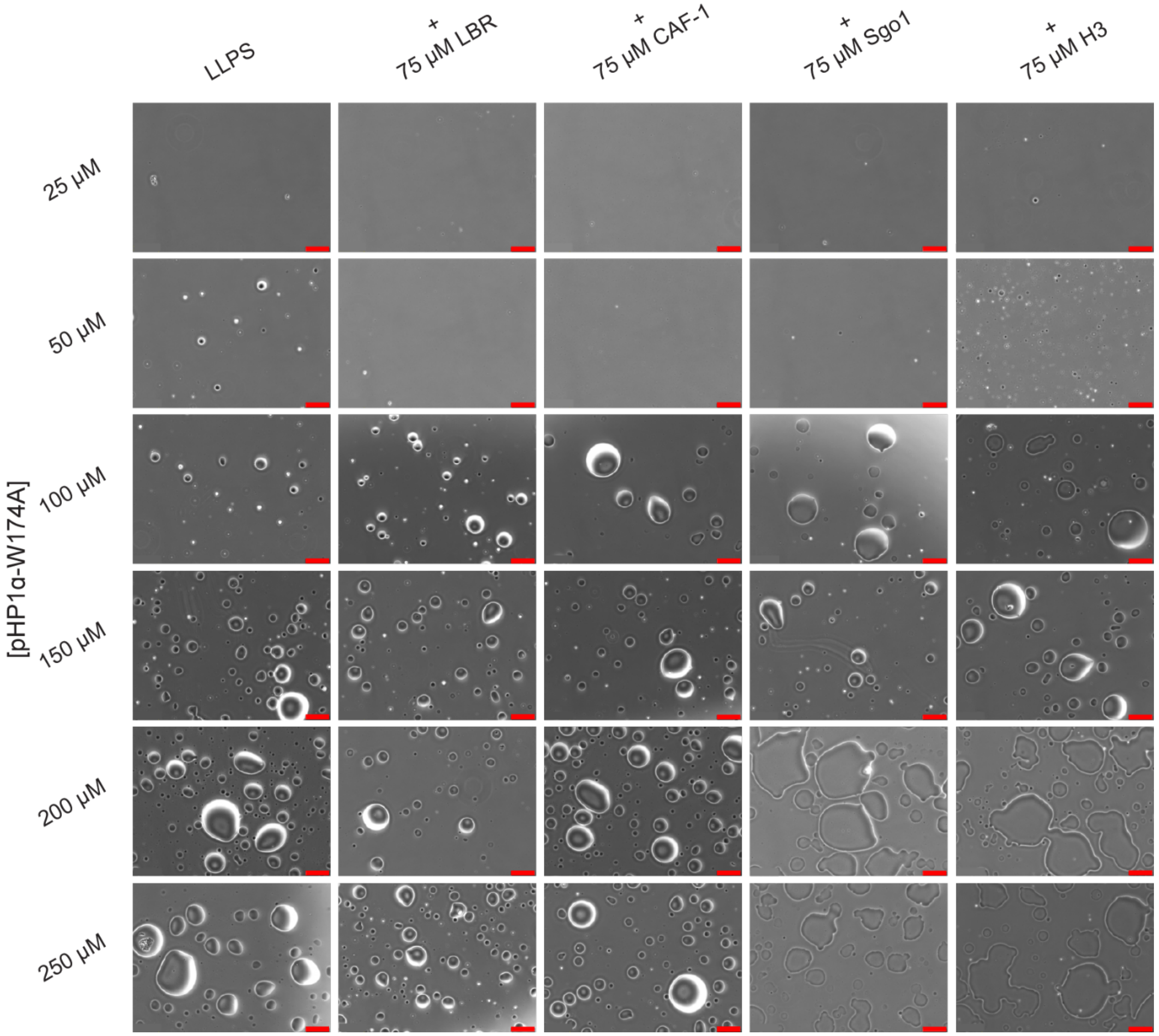


**Figure S13.** Brightfield microscopy images of LLPS of pHP1α-W174A LLPS with and without 75 μM peptide as indicated on top of each column. The red scale bar represents 100 μm.

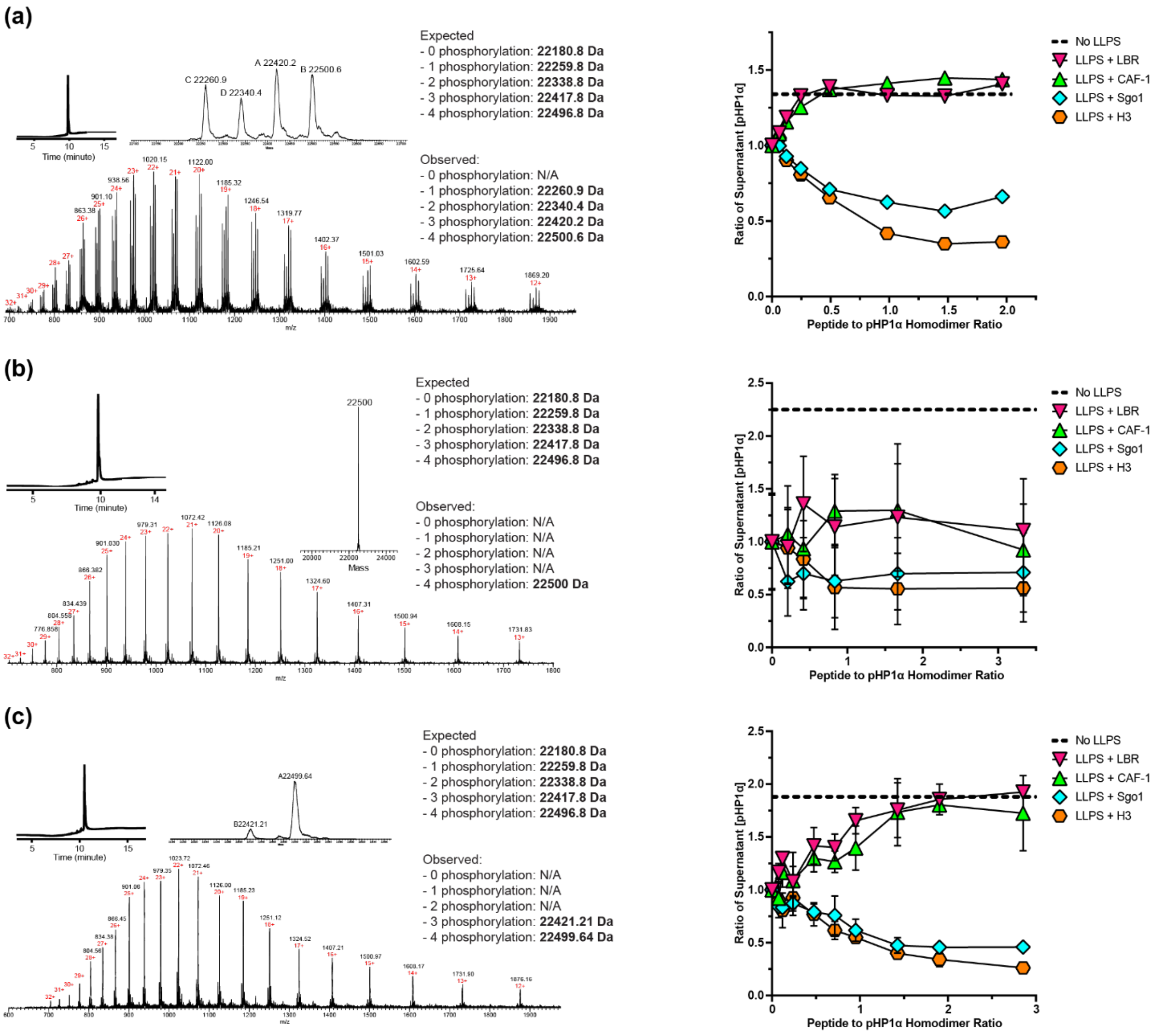


**Figure S14. (a) – (c)** LC-MS analysis of different phosphorylation batches of pHP1α. The plot of supernatant pHP1α concentration in the presence of the peptides showed consistent trend of the effect of each peptide on LLPS of pHP1α. Data from (a) were performed in one replicate. Experiments from (b) and (c) were performed in triplicate.

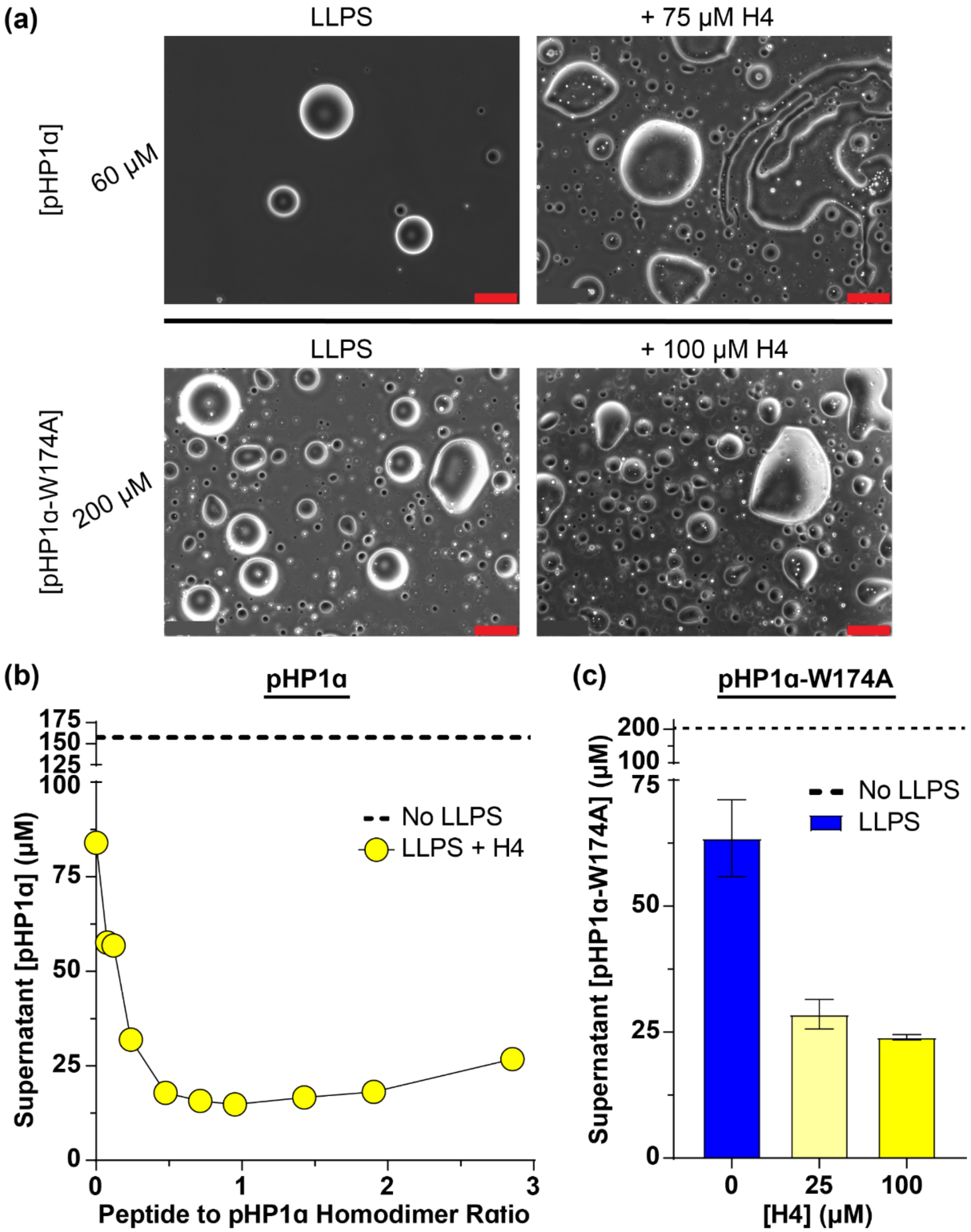


**Figure S15.** LLPS of pHP1α and pHP1α-W174A with and without H4 peptide. **(a)** Brightfield microscopy images of pHP1α and pHP1α-W174A with and without H4. The red scale bar represents 100 μm. LLPS supernatant concentration of **(b)** pHP1α, and **(c)** pHP1α-W174A with and without H4. The dashed line represents the expected supernatant concentration if LLPS does not occur. The error bars represent the standard deviation from measurements done in triplicate.

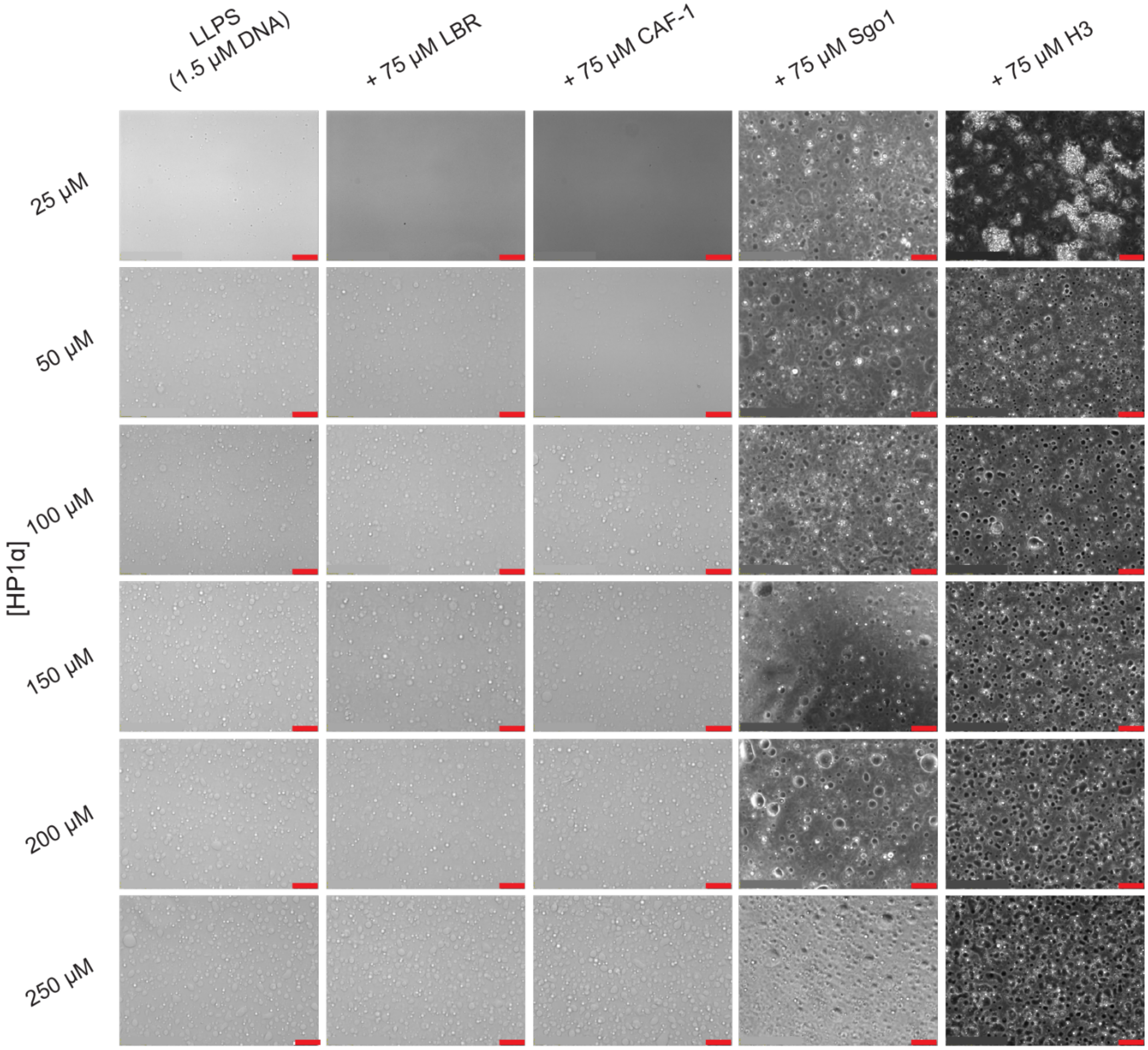


**Figure S16.** Brightfield microscopy images of HP1α LLPS using 1.5 μM of 205 bp DNA with and without 75 μM peptide as indicated on top of each column. The red scale bar represents 50 μm.

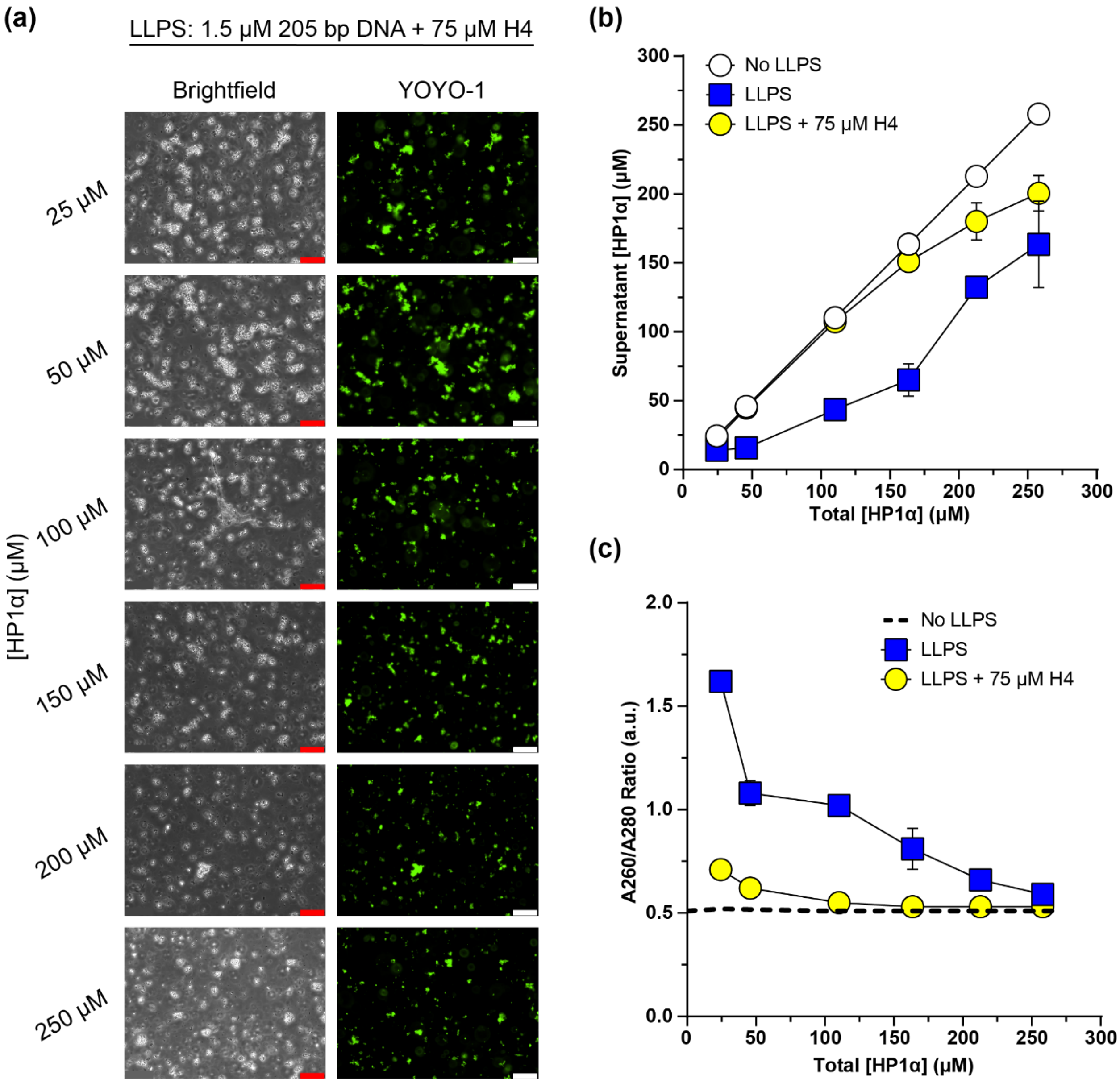


**Figure S17.** LLPS of HP1α using 205 bp DNA with and without H4 peptide. **(a)** Brightfield and fluorescence (using YOYO-1 dye) microscopy images. **(b)** Supernatant HP1α concentration after LLPS, and **(c)** A260/A280 ratio of HP1α LLPS with 75 μM H4 using varying concentrations of HP1α from 25 to 250 μM. The red and white scale bars represent 50 μm. The dashed line represents the expected supernatant concentration if LLPS does not occur and there is no DNA in the sample. Experiments were performed in triplicate.

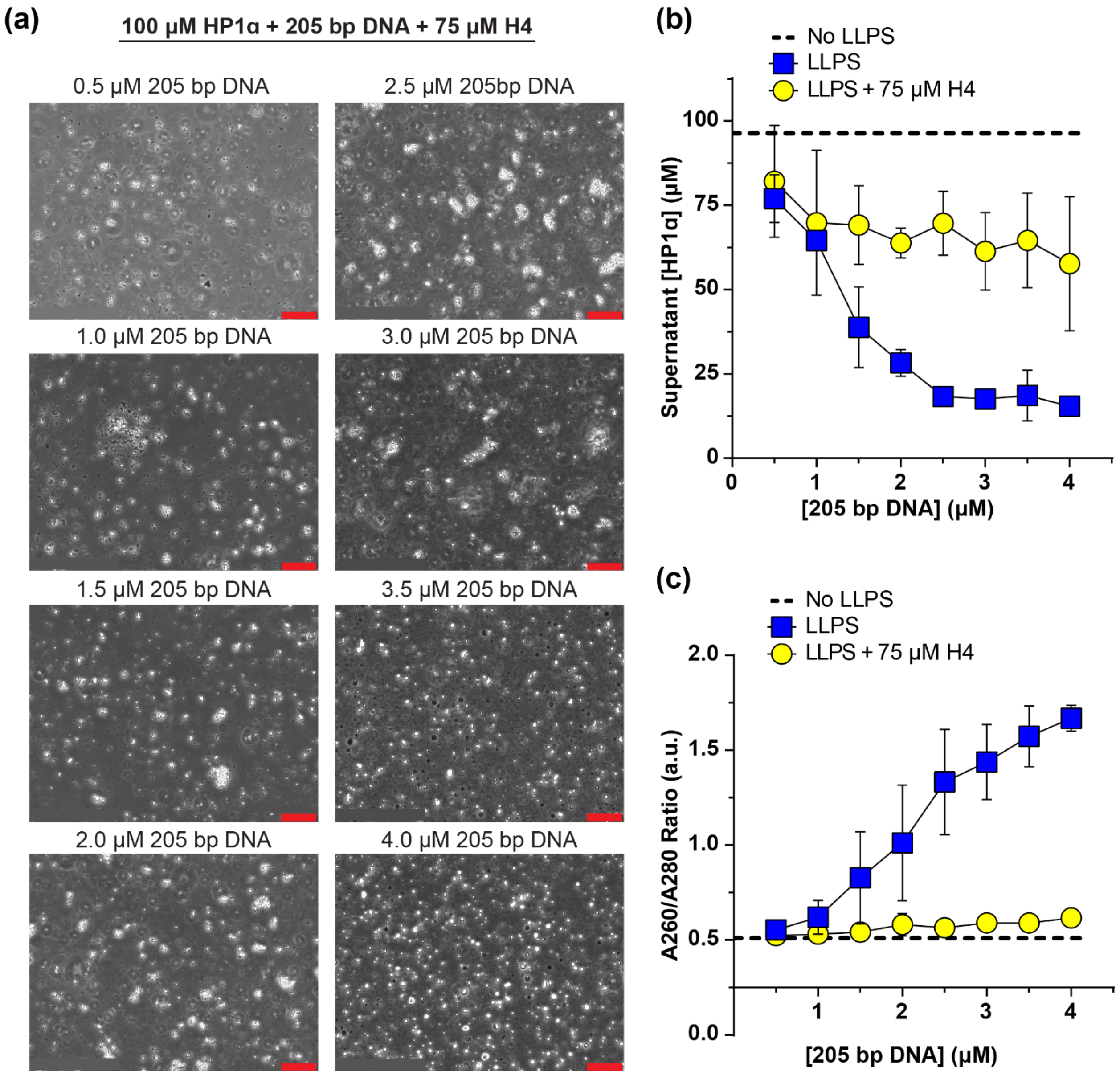


**Figure S18.** LLPS of HP1α using 205 bp DNA with and without H4 peptide. **(a)** Brightfield microscopy images, **(b)** supernatant HP1α concentration after LLPS, and **(c)** A260/A280 ratio of HP1α LLPS with 75 μM H4 using varying concentrations of DNA from 0.5 to 4.0 μM. The red scale bar represents 50 μm. The dashed line represents the expected supernatant concentration if LLPS does not occur and there is no DNA in the sample. Experiments were performed in triplicate.

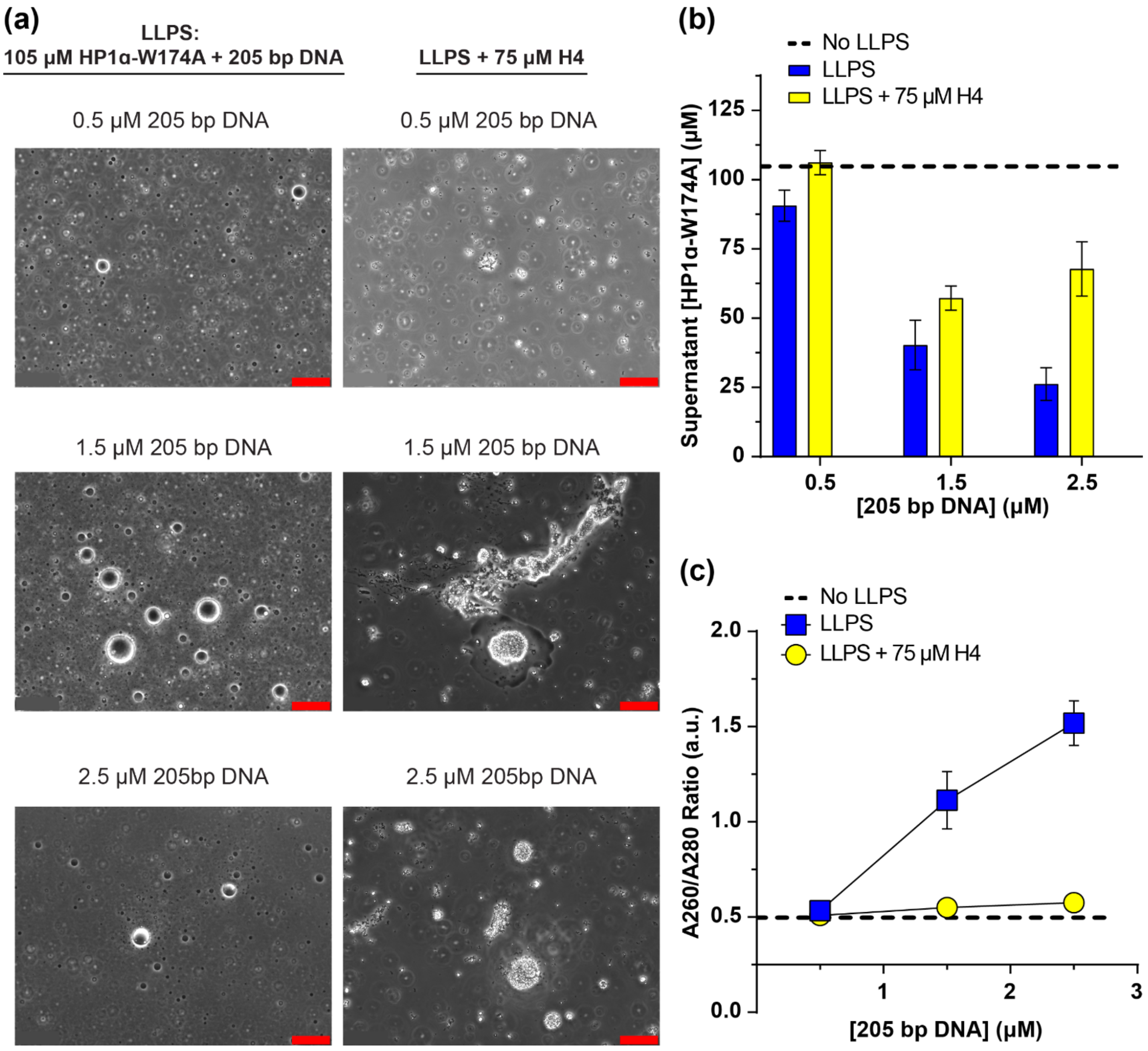


**Figure S19.** LLPS of HP1α-W174A using 205 bp DNA with and without H4. **(a)** Brightfield microscopy images, **(b)** supernatant HP1α-W174A concentration, and **(c)** A260/A280 ratio of HP1α-W174A LLPS with 75 μM H4 using varying concentrations of DNA from 0.5 to 4.0 μM. The red scale bar represents 50 μm. The dashed line represents the expected supernatant concentration if LLPS does not occur and there is no DNA in the sample. Experiments were performed in triplicate.

### Supporting information tables

**Table S1.** LC-MS/MS Analysis of pHP1α.

| Sequence (STY Phosphorylation Probabilities) | # of Phosphorylations | m/z (charge) [1] | Score | PEP [2] | Ratio Mod./Base [3] |
|---|---|---|---|---|---|
| RKS(0.983)NFS(0.017)NSADDIK | 1 | 520.231 (+3) | 93.181 | 2.61E-05 | 0.00006 |
| SNFSNS(1)ADDIK | 1 | 638.249 (+2) | 179.59 | 3.01E-75 | 0.00609 |
| T(0.015)ADS(0.977)S(0.91)S(0.549)S(0.549)EDEEEYVVEK | 3 | 1071.342 (+2) | 145.23 | 1.99E-47 | 0.63060 |
| T(0.073)ADS(0.934)S(0.924)S(0.924)S(0.14)EDEEEY(0.005)VVEK | 3 | 1071.342 (+2) | 145.23 | 1.99E-47 | 0.63060 |
| T(0.022)ADS(0.635)S(0.531)S(0.905)S(0.905)EDEEEY(0.002)VVEK | 3 | 1071.342 (+2) | 125.97 | 2.30E-39 | 0.10628 |
| LT(1)WHAYPEDAENK | 1 | 550.896 (+3) | 84.352 | 1.07E-05 | 0.00222 |
| WKDT(1)DEADLVLAK | 1 | 791.364 (+2); 527.576 (+3) | 172.32 | 2.29E-82 | 0.00012 |
| WKGFS(0.32)EEHNT(0.68)WEPEK | 1 | 660.940 (+3) | 54.292 | 0.00911895 | 0.00036 |
| LT(0.082)WHAY(0.918)PEDAENKEK | 1 | 636.608 (+3); 477.456 (+4) | 94.673 | 3.91E-07 | 0.00065 |

1) The mass to charge ratio of the modified peptide.
2) The posterior error probability (PEP), which is the probability of a peptide being a false hit.
3) The ratio of the modified peptide peak intensity over the base peak intensiity (unmodified peptide).

**Table S2.** Dissociation constants (Kd) of HP1α-CSD or full-length HP1α with different peptides as determined in the published literature.

| Peptide | Kd (μM) | Method | Reference |
|---|---|---|---|
| CAF-1* | 0.43 ± 0.10 | Tryptophan Fluorescence | Richart et al., J. Biol. Chem. (**2012**) |
| H3* | 58 ± 7 | NMR | Richart et al., J. Biol. Chem. (**2012**) |
| LBR** | 3 | Fluoresence Anisotropy | Larson et al., *Nature* (**2017**) |
| Sgo1** | 0.16 | Fluoresence Anisotropy | Larson et al., *Nature* (**2017**) |

* Kd determined with HP1α-CSD (residues 109-185).
** Kd determined with HP1α.

**Table S3.** Properties of the HP1α constructs and the peptide ligands.

| Protein/Peptide | Residues | ε at 280 nm (M-1cm-1)* | ε at 205 nm (M-1cm-1)** | Calculated Molecular Mass (Da)* | Source |
|---|---|---|---|---|---|
| HP1α | 1-191 | 29,160 | N/A | 22,181 | E. Coli Expression |
| HP1α-W174A | 1-191 (W174A) | 23,470 | N/A | 22,066 | E. Coli Expression |
| HP1α-I165E | 1-191 (I165E) | 29,160 | N/A | 22,197 | E. Coli Expression |
| HP1α-CSD | 112-176 | 12,660 | N/A | 7,487 | E. Coli Expression |
| HP1α-CSDCTE | 110-191 | 13,940 | N/A | 9,409 | E. Coli Expression |
| HP1α-CSDCTE-W174A | 110-191 (W174A) | 8,250 | N/A | 9,293 | E. Coli Expression |
| HP1α-CSDCTE-I165E | 110-191 (I165E) | 13,940 | N/A | 9,424 | E. Coli Expression |
| LBR | 105-124 | 0 | 55520 | 2,348 | Commercial Source |
| CAF-1 | 210-238 | 0 | 89220 | 3,111 | Commercial Source |
| Sgo1 | 446-466 | 1,280 | N/A | 2,440 | Commercial Source |
| H3 | 37-59 | 2,560 | N/A | 2,841 | Commercial Source |
| H4 | 1-24 | 0 | 74540 | 2,476 | Commercial Source |

* Values were calculated using the website *http://protcalc.sourceforge.net/* .
** Values were calculated using the website *https://spin.niddk.nih.gov/clore/* .
N/A — value not calculated.
