## Supplementary figures and images for "Molecular interactions underlying the phase separation of HP1α: Role of phosphorylation, ligand and nucleic acid binding"

### Movie S3

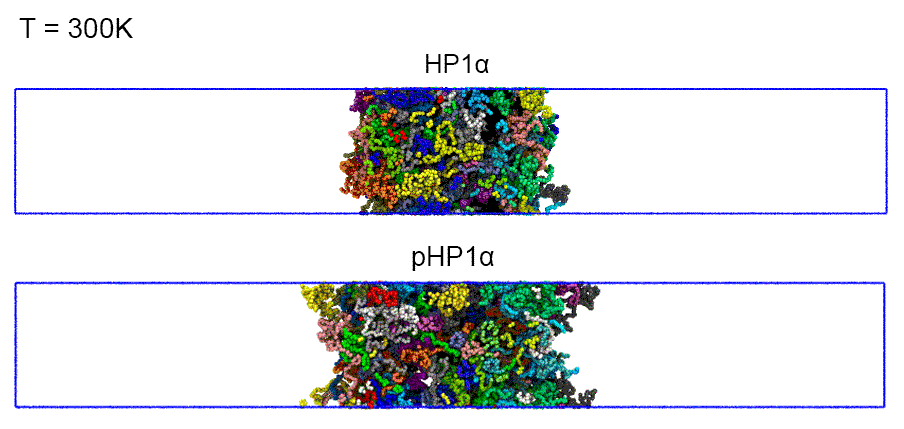

### Movie S4

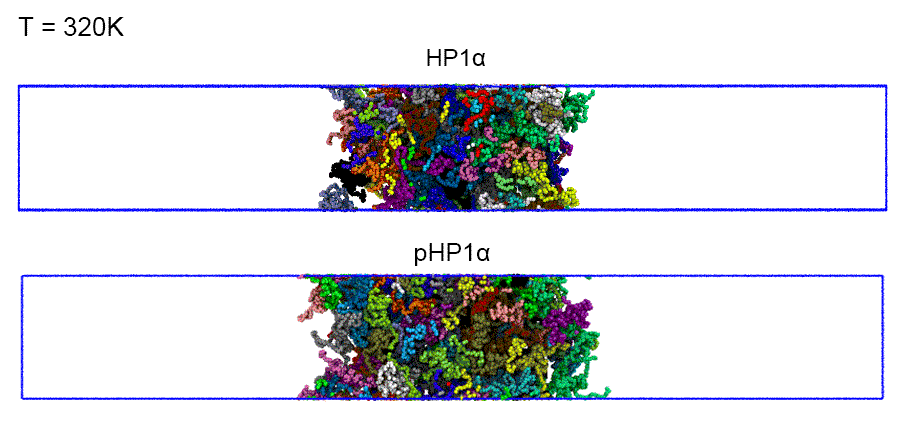

### Movie S5

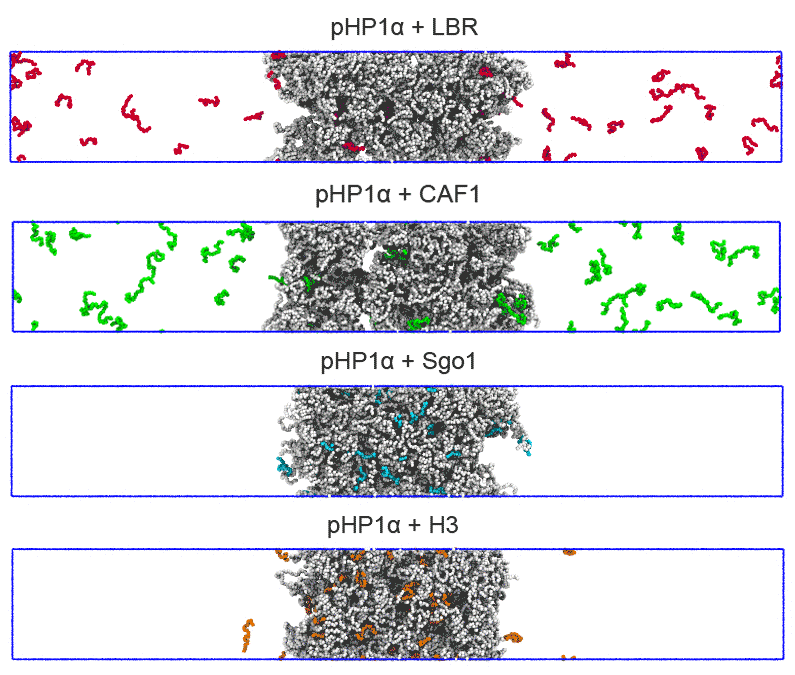

### Movie S6

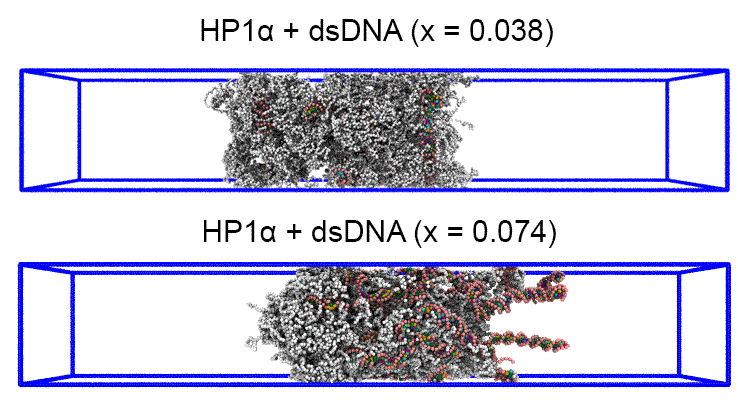
