## Supplementary material for "Molecular interactions underlying the phase separation of HP1α: Role of phosphorylation, ligand and nucleic acid binding": Movie captions

**Movie S1.** All-atom simulation of HP1α. The CSD-CSD domains were aligned and fixed for visualization.

**Movie S2**. All-atom simulation of pHP1α. The CSD-CSD domains were aligned and fixed for visualization. Phosphorylated serine residues are shown in licorice representation.

**Movie S3**. Slab simulations of HP1α and pHP1α homodimers at 300K.

**Movie S4**. Slab simulations of HP1α and pHP1α homodimers at 320K.

**Movie S5**. Slab simulations of pHP1α homodimers with peptides at a 1:1 peptide to pHP1α ratio at 320K.

**Movie S6**. Slab simulations of HP1α homodimers with different mole fractions of dsDNA at 320K.
